# Shoot nitrate status regulates Arabidopsis shoot growth and systemic transcriptional responses via shoot adenosine phosphate-isopentenyltransferase 3

**DOI:** 10.1101/2024.12.26.630360

**Authors:** Kota Monden, Takamasa Suzuki, Mikiko Kojima, Yumiko Takebayashi, Daisuke Sugiura, Tsuyoshi Nakagawa, Hitoshi Sakakibara, Takushi Hachiya

## Abstract

Plants systemically regulate growth and gene expression according to their internal nitrate status. Our previous study reported that shoot nitrate accumulation increases shoot expression of adenosine phosphate-*ISOPENTENYLTRANSFERASE 3* (*IPT3*) and shoot levels of *N*^6^-(Δ^2^-isopentenyl) adenine (iP)-type cytokinins (CKs). *IPT3* expression is localized in the phloem, and iP-type CKs, which are synthesized by IPT3, are phloem-mobile. As CKs are a class of phytohormones that control growth and genome-wide gene expression, shoot-expressed IPT3 may mediate the systemic regulation of growth and transcriptomic responses to shoot nitrate status. To examine this, we developed a novel system to manipulate nitrate levels and *IPT3* expression in a shoot-specific manner and performed growth analysis, CK determination, and RNA-sequencing. Our results demonstrated that shoot nitrate accumulation significantly promoted shoot growth and elevated shoot concentrations of iP ribotides, iP-7-*N*-glucoside, and iP-9-*N*-glucoside through the action of shoot IPT3. Transcriptomic responses to shoot nitrate accumulation were largely tuned by shoot IPT3, with opposite effects in shoots and roots. Shoot IPT3 amplified shoot responses of nitrate-inducible genes and immune response genes to shoot nitrate accumulation, while it dampened root responses of nitrate transport/assimilation genes. This transcriptomic modulation via shoot IPT3 was accompanied by coherent transcriptional changes in genes encoding mobile peptides and transcriptional repressors. Here we present a novel scheme integrating shoot nitrate status responses with CK signaling.

## Introduction

Land plants use soil nitrate and ammonium as primary nitrogen sources (Hachiya and Sakakibara 2017). Root-absorbed nitrate is reduced to ammonium via the sequential actions of nitrate reductase (NR) and nitrite reductase (NiR). Ammonium, whether taken up by the roots or generated through nitrate reduction, is assimilated into Gln and Glu by the enzymes glutamine synthetase (GS) and glutamate synthase (GOGAT). These amino acids serve as precursors for the biosynthesis of other amino acids, thereby providing carbon skeletons and amino groups for primary and secondary metabolism.

Nitrate serves as a building block for organic nitrogen biosynthesis and as a signaling molecule (Wang et al. 2004; Remans et al. 2006). The addition of exogenous nitrate to nitrate-depleted plants rapidly changes the expression of nitrate-responsive genes, a phenomenon termed the primary nitrate response (PNR) (Medici and Krouk 2014). In *Arabidopsis*, the nitrate transceptor Nitrate transporter 1.1 (NRT1.1) and the transcription-activating nitrate sensor NIN-like protein 7 (NLP7) play central roles in PNR (Ho et al. 2009; Bouguyon et al. 2015; Liu et al. 2017; Liu et al. 2022). NRT1.1 physically interacts with the Cyclic nucleotide-gated channel 15 to form a nitrate-activated Ca^2+^-permeable channel on the plasma membrane (Wang et al. 2021). Consequently, nitrate uptake via NRT1.1 transiently induces a Ca^2+^ influx, thereby elevating both cytosolic and nuclear Ca^2+^ levels (Riveras et al. 2015; Liu et al. 2017; Wang et al. 2021). Additionally, Phospholipase C also contributes to the NRT1.1-mediated increase in cytoplasmic Ca^2+^ levels (Riveras et al. 2015). The nitrate-binding affinity and uptake activity of NRT1.1 are tuned by specific protein kinases and phosphatase, including CBL-interacting protein kinases 8/23, Sucrose non-fermenting1-related protein kinase (SnRK) 2s, and ABA insensitive 2 (Ho et al. 2009; Hu et al. 2009; Léran et al. 2015; Su et al. 2020). The subgroup III Calcium-dependent protein kinases 10/30/32 (CPK10/30/32) perceive Ca^2+^ and phosphorylate NLP7 at S205, causing its nuclear retention by inhibiting nuclear export (Marchive et al. 2013; Liu et al. 2017). Nitrate-bound NLP7, along with transcription factors NLP2/6 and Nitrate regulatory gene 2—which physically interact with NLP7—promotes the transcription of PNR genes including those encoding secondary transcription factors, thereby further altering the expression of downstream targets (Konishi and Yanagisawa 2013; Xu et al. 2016; Maeda et al. 2018; Liu et al. 2022; Durand et al. 2023).

Besides its role in PNR, nitrate acts as an internal signal of plant nitrogen status, modulating nitrogen responses under steady-state conditions. In tobacco plants with different NR activities grown under varying nitrate availability, the shoot-to-root dry weight ratio was positively correlated with foliar nitrate levels (Scheible et al. 1997). In *Arabidopsis*, abundant nitrate supply inhibits primary and lateral root growth (Zhang et al. 1998; Signora et al. 2001). Patterson et al. (2016) reported that nitrate application increases the shoot expression of glutaredoxin (Grx) genes *GrxS3/4/5/8* via Arabidopsis response regulators (ARRs) 1/10/12 for cytokinin (CK) signaling, thereby inhibiting primary root growth. Kobayashi et al. (2024) suggested that GrxS1–8 are candidate phloem-mobile signals from nitrate-supplied shoots to roots, and they bind to TGACG-binding factor 1 (TGA1) and TGA4 (Alvarez et al. 2014) in roots, forming a corepressor complex with TOPLESS to limit high-affinity nitrate uptake. Meanwhile, Ota et al. (2020) discovered that under reduced shoot nitrogen, including nitrate depletion, C-terminally encoded peptide downstream-like 2 (CEPDL2)—a Grx family protein also known as GrxS9—is induced in the leaf vasculature and subsequently translocated to the roots. The co-activator complex composed of CEPDL2, its homologs CEPD1/2, and TGA1/4 promotes root nitrate uptake (Kobayashi et al. 2024). These findings imply that shoot nitrate status influences root nitrogen acquisition.

In our previous study, to focus on the action of shoot nitrate status, we manipulated shoot nitrate levels using wild-type *Arabidopsis* and an NR-null mutant (Okamoto et al. 2019). Subsequent RNA-sequencing (RNA-seq) suggested that shoot nitrate accumulation alone represses the expression of nitrogen starvation-inducible genes in shoots and roots. Notably, we observed that shoot nitrate accumulation was accompanied by increases in the shoot expression of adenosine phosphate-*ISOPENTENYLTRANSFERASE 3* (*IPT3*) and the shoot levels of *N*^6^-(Δ^2^-isopentenyl) adenine (iP)-type CKs, including iP ribotides (iPRPs), iP-7-*N*-glucoside (iP7G), and iP-9-*N*-glucoside (iP9G) (Hachiya et al. 2020). Adenosine phosphate-IPTs including IPT3 are key enzymes that catalyze the formation of iPRPs using DMAPP as the isoprenoid donor and adenosine phosphates as acceptors (Sakakibara 2021). iP riboside 5′-monophosphate (iPRMP) is then activated to bioactive free base iP via either the one-step Lonely guy (LOG) pathway or a two-step pathway involving 5′-ribonucleotide phosphohydrolase GY3 and CK/purine riboside nucleosidase CPN1/NSH3 (Kurakawa et al. 2007; Kojima et al. 2023; Wu et al. 2023). iP homeostasis is further fine-tuned through *N*-glucosylation catalyzed by glucosyltransferases (UGTs) and degradation by cytokinin oxidase/dehydrogenases (CKXs) (Werner et al. 2003; Hošek et al. 2020). CKs are a class of phytohormones that regulate a wide range of developmental and physiological processes, including cell division, shoot and root growth, leaf senescence, vascular formation, and responses to biotic and abiotic stresses (Werner and Schmülling 2009; Cortleven et al. 2019). Multiple transcriptome analyses have demonstrated that CKs regulate specific gene sets across several tissues and organs (Bhargava et al. 2013). Nitrate-induced expression of *IPT3* is localized mainly in phloem cells (Miyawaki et al. 2004; Takei et al. 2004a), and iP-type CKs, which are de novo synthesized by IPT3, are phloem-mobile (Hirose et al. 2008; Bishopp et al. 2011). Hence, the systemic regulation of growth and transcriptomic responses to shoot nitrate status may be mediated via IPT3-synthesized iP-type CKs. Supporting this idea, several lines of evidence indicate both overlap and crosstalk between nitrate and CK signaling. For instance, nitrate- and CK-responsive genes have been found to partially overlap (Sakakibara et al. 2006). In addition, the nitrogen depletion-dependent induction of shoot *CEPDL2* is enhanced by CKs translocated from roots (Ota et al. 2020). Furthermore, *trans*-zeatin (*t*Z)-type CKs mediate preferential root growth in nitrate-rich patches over nitrate-poor patches (Ruffel et al. 2011; Poitout et al. 2018).

Based on this context, the present study examined the systemic responses to shoot nitrate status and their dependence on shoot IPT3. We developed a novel experimental system to manipulate nitrate levels and *IPT3* expression in a shoot-specific manner using grafted plants derived from plants lacking NR and/or IPT3. Using the shoots and roots of the grafted plants, growth analysis, CK species determination, and RNA-seq were performed. Our results indicated that shoot nitrate accumulation enhances shoot growth and alters the transcriptomes of shoots and roots, which is elaborately tuned by shoot IPT3 in the opposite direction in shoots and roots. We herein present a scheme incorporating shoot nitrate accumulation responses with shoot IPT3.

## Materials and Methods

### Plant materials

In this study, we used *A. thaliana* accession Columbia (Col) as the control line; NR-null mutant (NR-null, Wang et al. 2007), *IPT3*-knockout mutant (*ipt3*, Miyawaki et al. 2006), and *IPT3/IPT5/IPT7*-knockout mutant (*ipt3,5,7*, Miyawaki et al. 2006); and a transgenic plant expressing *GFP* driven by the *IPT3* promoter (*proIPT3:GFP*, Takei et al. 2004a). Homozygous multiple mutants of NR-null *ipt3* and NR-null *proIPT3:GFP* were produced via crossing.

### Culture media

Three different solid media were used for *in vitro* plant culture as follows: (i) 2.5 mM NH_4_^+^ medium, half-strength Murashige and Skoog (1/2MS) modified basal salt mixture without nitrogen (M531; PhytoTech Labs, Lenexa, KS, USA) supplemented with 1.25 mM (NH_4_)_2_SO_4_, 10.0 g L^−1^ sucrose, 1.0 g L^−1^ MES, and 4.0 g L^−1^ gellan gum (Fujifilm Wako, Osaka, Japan) and adjusted to pH 6.7 with KOH; (ii) 10 mM NO_3_^−^ and 2.5 mM NH_4_^+^ medium, 1/2MS modified basal salt mixture without nitrogen (M531; PhytoTech Labs) supplemented with 10 mM KNO_3_, 1.25 mM (NH_4_)_2_SO_4_, 10.0 g L^−1^ sucrose, 1.0 g L^−1^ MES, and 4.0 g L^−1^ gellan gum (Fujifilm Wako) and adjusted to pH 6.7 with KOH; and (iii) no nitrogen medium, 1/2MS modified basal salt mixture without nitrogen (M531; PhytoTech Labs) supplemented with 2.0 g L^−1^ sucrose, 0.5 g L^−1^ MES, and 5.0 g L^−1^ gellan gum (Fujifilm Wako) and adjusted to pH 5.7 with KOH. It should be noted that pH 6.7 was employed to mitigate ammonium toxicity under 2.5 mM NH_4_^+^ medium (Hachiya et al. 2021b) and to promote better growth of NR-deficient plants under 10 mM NO_3_^−^ and 2.5 mM NH_4_^+^ medium (Wang et al. 2004).

### Transfer experiments using grafted plants

To manipulate nitrate levels and *IPT3* expression in a shoot-specific manner, micrografting and transfer experiments were performed using Col, NR-null, *ipt3*, NR-null *ipt3*, and *ipt3,5,7* plants (Figure 1A, B). Surface-sterilized seeds were stratified in the dark at 4°C for 3 days. To ensure uniform growth regardless of the NR activity, approximately 120 seeds per dish were sown in rectangular dishes (140 × 100 × 20 mm^3^; Eiken Chemical Co. Ltd., Tokyo, Japan) containing 50 mL of 2.5 mM NH_4_^+^ medium and grown vertically at 22°C for 4 days under a photosynthetic photon flux density (PPFD) of 25 µmol m^−2^ s^−1^ with a 16/8-h light/dark cycle (condition 1). Four-day-old seedlings were grafted aseptically using four scion/rootstock combinations: Col/*ipt3,5,7*, NR-null/*ipt3,5,7*, *ipt3*/*ipt3,5,7*, and NR-null *ipt3*/*ipt3,5,7* (Figure 1B). Notably, *ipt3,5,7* was used as the rootstock to avoid the complementation of shoot-specific CK depletion in *ipt3* by root-derived CKs. The seedlings were cut perpendicularly at the upper hypocotyl with an injection needle tip (NN-2613S; TERUMO, Tokyo, Japan) on a membrane filter (HAWP04700; Merck Millipore, Darmstadt, Germany). The obtained scion was connected with the partner rootstock through a 0.4-mm silicone microtube (1-8194-04; AS ONE, Osaka, Japan). The grafted plants were incubated at 27°C for 5 days under a PPFD of 50 µmol m^−2^ s^−1^ with a constant light (condition 2) to enhance grafting success (Monden et al. 2022), and further cultivated at 22°C for 24 h under a PPFD of 25 µmol m^−2^ s^−1^ with a 16/8-h light/dark cycle (condition 3). Successfully grafted plants without adventitious root formation were selected for size uniformity. The grafted plants were transferred to round dishes (diameter, 90 mm; depth, 20 mm; AS ONE) containing 30 mL of 10 mM NO_3_^−^ and 2.5 mM NH_4_^+^ medium and grown for 24 h under a PPFD of 25 µmol m^−2^ s^−1^ with a constant light to promote nitrate accumulation in the grafted plants (condition 4). Finally, the plants were transferred to round dishes containing 30 mL of no nitrogen medium and grown for 5 days under a PPFD of 25 µmol m^−2^ s^−1^ with a constant light (condition 5) to allow nitrate accumulation in the shoots of NR-null/*ipt3,5,7* and NR-null *ipt3*/*ipt3,5,7* but not in Col/*ipt3,5,7* and *ipt3*/*ipt3,5,7*. No external nitrogen such as ammonium was supplied to avoid enrichment of internal nitrogen pools and unintended signaling, which could obscure the effects of shoot nitrate status. The 5-day treatment was chosen to ensure sufficient growth and transcriptomic responses while minimizing transient stress caused by the pH drop from condition 4 to condition 5. In addition, a low light intensity of 25 µmol m^−2^ s^−1^ was employed to reduce stress, given that high light intensity is known to exacerbate nitrogen deficiency stress (Iwagami et al. 2022). The surfaces of no nitrogen medium were covered with cellophane sheets to allow nondestructive sampling of the roots (Hachiya et al. 2021a). In conditions 2–5, four types of grafted plants were grown under identical external nitrogen conditions to ensure uniform biosynthesis of systemic signaling molecules responsive to external nitrogen availability, such as C-terminally encoded peptides (CEPs) (Tabata et al. 2014), thereby isolating the systemic effects of shoot nitrate status and enabling clearer interpretation of the results.

**Figure 1.**
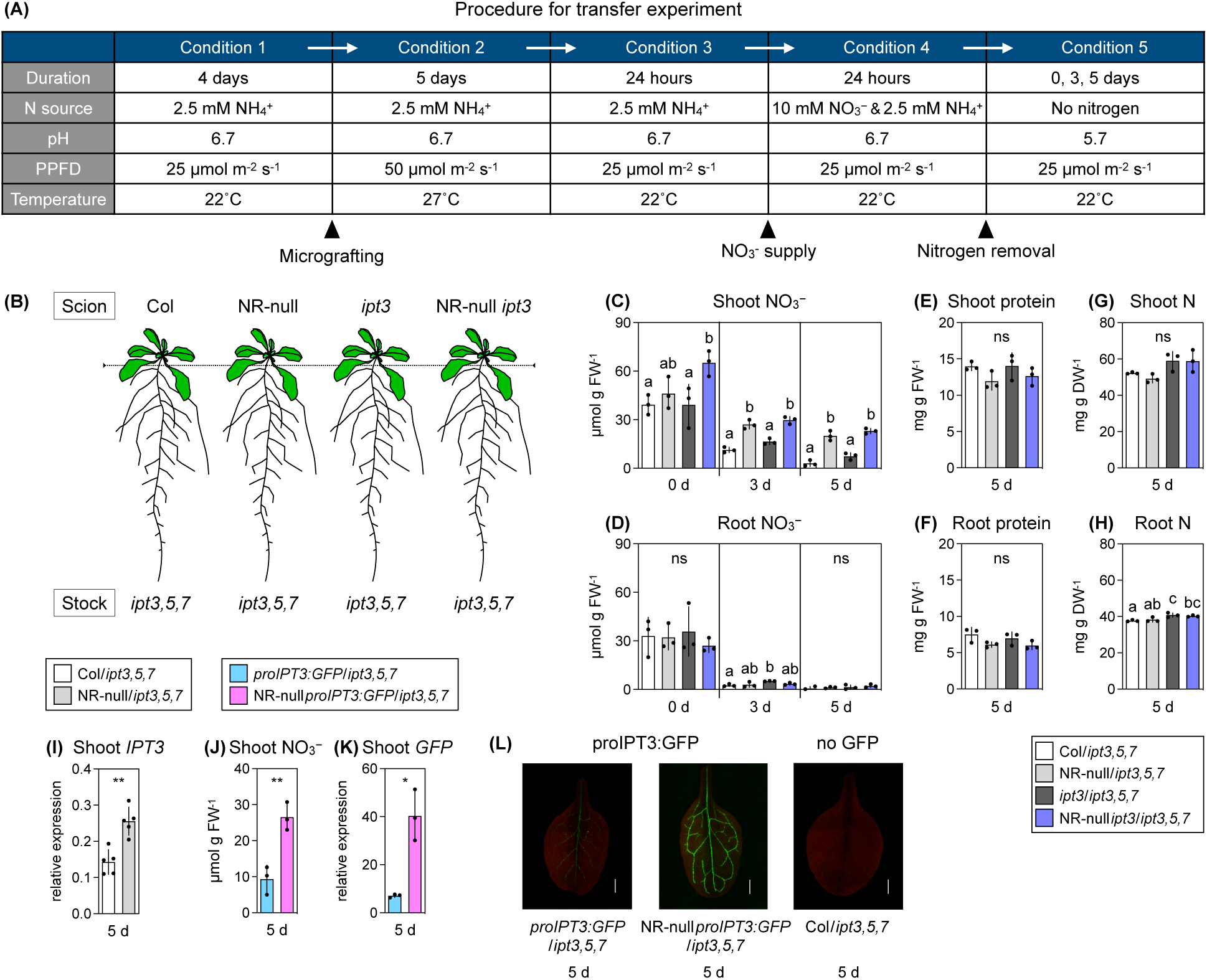
Micrografting and transfer experiments for manipulation of nitrate levels and *IPT3* expression in a shoot-specific manner. (A) Summary of the micrografting and transfer experiments. (B) A schematic representation of four types of grafted plants. The dotted line represents the grafted seam in the hypocotyl. (C, D) Nitrate concentrations in shoots (C) and roots (D) 0, 3, or 5 days after nitrogen removal. (E, F) Total protein concentrations in shoots (E) and roots (F) 5 days after nitrogen removal. (C–F) Two grafted plants were pooled as one biological replicate. Data are presented as mean ± SD (n = 3). (G, H) Total nitrogen concentrations in shoots (G) and roots (H) 5 days after nitrogen removal. Sampling was performed three times independently. In this sampling, 7, 6, and 5 grafted plants were pooled as one biological replicate. Data are presented as mean ± SD (n = 3). (C–H) Different lowercase letters indicate significant differences, as determined via Tukey–Kramer test (*P* < 0.05). (I) Transcript levels of *IPT3* in shoots 5 days after nitrogen removal, as determined using RT-qPCR. Sampling was done five times independently. In this sampling, 9, 6, 7, 7, and 8 grafted plants were pooled as one biological replicate. Data are presented as mean ± SD (n = 5). (J) Nitrate concentrations in shoots 5 days after nitrogen removal. (K) Transcript levels of *GFP* in shoots 5 days after nitrogen removal, as determined via RT-qPCR. (J, K) One biological replicate consisted of one grafted plant. Data are presented as mean ± SD (n = 3). (I–K) Statistical significance was assessed using unpaired two-tailed Welch’s *t*-test (\**P* < 0.05; \*\**P* < 0.01; \*\*\**P* < 0.001). (L) Fluorescence images of true leaves 5 days after nitrogen removal. Green and red colors correspond to GFP fluorescence and autofluorescence, respectively. Scale bars represent 500 μm. In graph legends, the grafted plants are indicated as “scion line name/rootstock line name.” ns, not significant; FW, fresh weight; DW, dry weight.

### Growth analysis

Shoots and roots were weighed on a precision balance (HR-202i; A&D, Tokyo, Japan), and shoots were scanned at a resolution of 600 dpi (GT-X980; EPSON, Tokyo, Japan) to measure the rosette diameter. Leaf blades were harvested from the shoots and scanned at a resolution of 600 dpi to measure the blade area of the cotyledon and true leaf. The rosette diameter and leaf blade area were measured using ImageJ version 1.52p (US National Institutes of Health, Bethesda, MD, USA).

### Determination of the total nitrogen content

The total nitrogen concentration was quantified as previously described (Monden et al. 2022). Harvested shoots and roots were oven-dried at 80°C for 2 days, and the dried samples were ground using a spoon and stored in a desiccator before use. The dried powder was weighed and encapsulated in a tin boat. Nitrogen concentrations were measured using a CN analyzer (Vario EL III; Elementar Analysensysteme GmbH, Hanau, Germany).

### Determination of the nitrate content

Nitrate content was quantified as previously described (Okamoto et al. 2019) with slight modifications. Frozen samples were ground with a Multi-Beads Shocker (Yasui Kikai, Osaka, Japan) using zirconia beads (diameter, 5 mm). The powder samples were boiled with 10 volumes of Milli-Q water at 100°C for 20 min. Five microliters of the supernatant or 5 µL of standard solution as dilution series of potassium nitrate was mixed with 40 µL of 0.05% (w/v) salicylic acid in sulfuric acid. The resulting mixture was then incubated at room temperature for 20 min. A mock treatment of 40 µL of sulfuric acid alone was added to 5 µL of the supernatant. One microliter of 8% (w/v) NaOH solution was added to the mixture, and the nitrate content was determined by measuring the absorbance at 410 nm using a plate reader (CORONA SH-9000Lab, Hitachi High-Tech Corp., Tokyo, Japan).

### Observation of fluorescent signals

Fluorescent imaging was performed using an All-in-one fluorescence microscope (BZ-X710, KEYENCE, Osaka, Japan) with a 4× objective (CFI Plan Fluor DL 4×/0.13, Nikon, Tokyo, Japan) and a metal halide lamp (No. 91056, KEYENCE). A narrow-band GFP filter (ex. 480/20; em. 510/20, 49020, Chroma, Brattleboro, VT, USA) and Cy5 filter (ex. 620/60; em. 700/75, 49006, Chroma) were used to detect the fluorescence of GFP and chlorophyll, respectively.

### Determination of CK species

The extraction and determination of CKs were performed with slight modifications to the method described by Kojima et al. (2009). Frozen tissue was ground into powder using a TissueLyser (Qiagen, Hilden, Germany) with a 5 mm zirconia bead. Samples were extracted in 1 ml of methanol : formic acid : water (15 : 1 : 4), with stable isotope-labeled internal standards. After incubation at –30°C for ≥16 h, samples were centrifuged at 10,000 × *g*, for 15 min. The supernatant was collected, and the pellet was re-extracted with 0.2 ml of the same solvent. Combined extracts were purified using Oasis HLB 96-well plates (Waters, Milford, MA, USA) on a STAR (HAMILTON, Reno, NV, USA), followed by evaporation and reconstitution in 1 ml of 1 M formic acid. Fractionation was performed on an Oasis MCX 96-well plate (Waters). ABA, auxins, and gibberellins were eluted with methanol (Elution 1); CK nucleotides with 0.35 M ammonia (Elution 2); and CK nucleobases, nucleosides, and glucosides with 0.35 M ammonia in 60% methanol (Elution 3). All fractions were dried by evaporation. For CK analysis, Elution 3 was reconstituted in 50 μl of 0.1% acetic acid. Elution 2 was reconstituted in 0.84 ml of 0.1 M CHES-NaOH (pH 9.8), treated with 1 U µl^-1^ of alkaline phosphatase (Oriental Yeast, Tokyo, Japan), incubated at 37°C for 1 h, neutralized, desalted via Oasis HLB plate, evaporated, and reconstituted in 50 μl of 0.1% acetic acid. CKs were analyzed using an ultra-performance liquid chromatography–tandem quadrupole mass spectrometry system (ACQUITY UPLC System/XEVO-TQS; Waters) with an octadecylsilyl column (ACQUITY UPLC HSS T3, 1.8 µm, 2.1 mm × 100 mm; Waters). CK species were separated at a flow rate of 0.25 mL min-1 with linear gradients of solvent A (0.06% acetic acid) and solvent B (0.06% acetic acid in methanol) set according to the following profile: 0 min, 95% A + 5% B; 7min, 60% A + 40% B; 10 min, 1% A + 99% B; 10 min, 1% A + 99% B; 13 min, 95% A + 5% B.

### Determination of the total protein content

The total protein content was determined as previously described (Hachiya et al. 2016). Frozen samples were ground with a Multi-Beads Shocker using zirconia beads (diameter, 5 mm). Total proteins were extracted with 10 vol. of Laemmli buffer (2% (w/v) SDS, 62.5 mM Tris-HCl (pH 6.8), 10% (v/v) glycerol, and 0.0125% (w/v) bromophenol blue) containing Halt™ protease inhibitor cocktail (Thermo Fisher Scientific, Waltman, MA, USA), followed by incubation at 95°C for 5 min. Extracts were centrifuged at 20,400 × *g* at room temperature for 10 min, and 10 μL of each aliquot was suspended in 500 μL of deionized water. Next, 100 μL of 0.15% (w/v) sodium deoxycholate was added, and the mixture was incubated at room temperature for 10 min. One hundred microliters of 72% (v/v) trichloroacetic acid was added, followed by incubation at room temperature for 15 min and centrifugation at 20,400 × *g* for 10 min. The precipitates were air-dried and suspended in 25 µL of deionized water. The suspension was analyzed with a TaKaRa BCA Protein Assay Kit (T9300A, TaKaRa, Kusatsu, Japan) according to the manufacturer’s instructions. Protein content was determined by measuring the absorbance at 562 nm using a plate reader (Hitachi High-Tech Corp.).

### RNA extraction

Frozen samples were ground with a Multi-Beads Shocker using zirconia beads (diameter, 5 mm). Total RNA was extracted using an RNeasy Plant Mini Kit (Qiagen) according to the manufacturer’s instructions. For RNA-seq library preparation, RNA was purified via on-column DNase digestion (Qiagen).

### RT-qPCR and RT-sqPCR

Reverse transcription (RT) was performed using a ReverTraAce qPCR RT Master Mix with gDNA Remover (Toyobo, Osaka Japan). RNA solution containing 0.5 µg of template RNA served as the input. Synthesized cDNA was then diluted 5-fold with water and used for quantitative PCR (qPCR) and semi-quantitative PCR (sqPCR). RT-qPCR was performed on a QuantStudio 1 (Thermo Fisher Scientific) with KOD SYBR qPCR Mix (Toyobo). Relative transcript levels were calculated using the comparative cycle threshold method, with *ACTIN3* as the internal standard (Hachiya et al. 2021b). RT-sqPCR was done on a T100 Thermal Cycler (Bio-Rad Laboratories, Tokyo, Japan) with Quick Taq HS (Toyobo) under appropriate cycling conditions. Primer sequences are presented in Table S1.

### RNA-seq

RNA quality was evaluated using a Qubit RNA IQ Assay Kit with a Qubit 4 Fluorometer (Thermo Fisher Scientific). The RNA samples for which the RNA IQs ranged 9.4–10.0 were used for RNA-seq. cDNA libraries were constructed using a NEBNext Ultra II RNA Library Prep with Sample Purification Beads (E7775S, New England Biolabs, Ipswich, MA, USA), a NEBNext Poly(A) mRNA Magnetic Isolation Module (E7490S, New England Biolabs), and a NEBNext Multiplex Oligos for Illumina (E7710S, New England Biolabs) according to the manufacturer’s instructions. These cDNA libraries were sequenced using NextSeq 500 (Illumina, San Diego, CA, USA), and the produced bcl files were converted to fastq files using bcl2fastq (Illumina). The reads were analyzed according to a previously described method (Notaguchi et al. 2014) and mapped to the Arabidopsis reference (TAIR10) using Bowtie (Langmead et al. 2009) with the following options: “--all --best --strata.” The mapped reads were processed using iDEP version 0.96 and 2.01 (Ge et al. 2018). For data normalization, the read counts data were transformed with rlog and normalized by subtracting the average expression for each gene according to the default setting of iDEP. For DEG extraction, DESeq2 was run with a false-discovery rate cutoff of 0.1. The enrichment analysis of each gene set was conducted using Metascape (Zhou et al. 2019) according to the default setting. Detailed information derived from the RNA-seq data is presented in Table S2–S9.

### Statistical analysis

Unpaired two-tailed Welch’s *t*-test, Tukey–Kramer multiple comparison test, two-way analysis of variance (ANOVA), and hypergeometric test were performed using R software v.2.15.3 (R Foundation for Statistical Computing, Vienna, Austria). The ANOVA tables are presented in Table S10.

## Results and Discussion

### Development of a novel system that manipulates nitrate levels and *IPT3* expression in a shoot-specific manner

To manipulate nitrate levels and *IPT3* expression in a shoot-specific manner, micrografting and transfer experiments were performed using Col, NR-null, *ipt3*, NR-null *ipt3*, and *ipt3,5,7* plants (Figure 1A, B). The plants were grown for 4 days in 2.5 mM NH_4_^+^ medium to ensure uniform growth regardless of the NR activity (condition 1). Then, grafted plants corresponding to Col/*ipt3,5,7*, NR-null/*ipt3,5,7*, *ipt3*/*ipt3,5,7*, and NR-null *ipt3*/*ipt3,5,7* (scion/rootstock) were generated (conditions 2 and 3). Notably, *ipt3,5,7* was used as the rootstock to avoid the complementation of shoot-specific CK depletion in *ipt3* by root-derived CKs. RT-sqPCR with primers specific for *NIA1*, *NIA2*, *IPT3*, *IPT5*, *IPT7*, and *ACTIN2* confirmed the successful generation of the desired grafted plants (Figure S1A, B). The grafted plants were transferred to 10 mM NO_3_^−^ and 2.5 mM NH_4_^+^ medium and incubated for 24 h to allow nitrate accumulation (condition 4). Then, the plants were transferred to no nitrogen medium and grown for 5 days (condition 5). After nitrogen removal, nitrate concentrations generally decreased over time in the shoots and roots of all plants (Figure 1C, D). At 3 and 5 days after nitrogen removal, the shoot nitrate concentrations were higher in NR-null/*ipt3,5,7* and NR-null *ipt3*/*ipt3,5,7* plants than in the other plants (Figure 1C), whereas root nitrate was depleted to the same level in all plants (Figure 1D). Meanwhile, at 5 days after nitrogen removal, total protein and total nitrogen levels in shoots and roots differed little among the plants (Figure 1E–H). RT-qPCR and RNA-seq revealed that shoot *IPT3* expression was significantly higher in NR-null/*ipt3,5,7* than in Col/*ipt3,5,7* (Figure 1I and Figure S1C). Only minor differences in transcripts per million (TPM) of shoot *IPT1/4/5/6/7/8* and root *IPT1/4/6/8* were observed among the four grafted plants (Figure S1C), suggesting that adenosine phosphate-*IPT* genes other than *IPT3* did not compensate for the absence of shoot IPT3. To visualize the tissue-specific induction of *IPT3* as a result of shoot NR deficiency, scions from *proIPT3:GFP* and NR-null *proIPT3:GFP* plants were grafted onto *ipt3,5,7* rootstocks, followed by the same transfer experiment as before. Shoot nitrate accumulation and *GFP* induction were observed in NR-null *proIPT3:GFP*/*ipt3,5,7* plants 5 days after nitrogen removal (Figure 1J, K), with strong GFP fluorescence localized in the leaf vein (Figure 1L). These findings suggest that *IPT3* induction due to shoot nitrate accumulation occurs mainly in the vasculature. Given that NLP7 promotes the *IPT3* transcription in response to nitrate application (Liu et al. 2022), it is plausible that *IPT3* induction in NR-null/*ipt3,5,7* is also mediated by NLP7. Overall, we successfully developed a novel system that manipulates nitrate levels and *IPT3* expression in a shoot-specific manner, with minimal impact on the total protein and total nitrogen levels within the plant, in the absence of external nitrogen sources. This system allows the analysis of systemic responses to shoot nitrate status and their dependence on shoot IPT3 in the following experiments. Furthermore, we confirmed that the TPM of nitrogen starvation-inducible *CEPs* (Tabata et al. 2014) did not differ significantly among the roots of grafted plants (Figure S1D). This enables us to isolate the systemic effects of shoot nitrate status and shoot IPT3 independently of root-derived CEP systemic signaling. All experiments described below were conducted using samples collected 5 days after nitrogen removal (condition 5).

### Shoot nitrate accumulation elevates shoot levels of iP-type CKs via shoot IPT3

To assess the impact of shoot nitrate status and shoot *IPT3* expression on CK activity, we measured the concentrations of iP-type and *t*Z-type CKs. The experiments were performed using the shoots and roots of grafted plants, with three biological replicates collected independently (Figure 2) or simultaneously (Figure S2). The levels of iP, iPRPs, iP7G, iP9G, *t*Z, *t*Z ribotides (*t*ZRPs), *t*Z-7-*N*-glucoside (*t*Z7G), *t*Z-9-*N*-glucoside (*t*Z9G), *t*Z-*O*-glucoside (*t*ZOG), and *t*ZR-*O*-glucoside (*t*ZROG) were consistently lower in the shoots of *ipt3*/*ipt3,5,7* and NR-null *ipt3*/*ipt3,5,7* plants than in those of Col/*ipt3,5,7* and NR-null/*ipt3,5,7* plants (Figure 2A, C–F, 2H–L and Figure S2A, C–F, S2H–L). Meanwhile, little difference was observed in the root levels of CK species among the grafted plants. These observations indicated that shoot IPT3 is crucial for maintaining the shoot levels of *t*Z-type and iP-type CKs, but it has only a minor effect on root CK levels. Notably, the shoot levels of iPRPs, iP7G, and iP9G were significantly increased by shoot NR deficiency in the presence of shoot IPT3 but not in its absence (Figure 2C–E and Figure S2C–E). This finding suggests that shoot nitrate accumulation elevates the shoot levels of these iP-type CKs via shoot IPT3. However, no significant increase was observed in the levels of iP, the biologically active form (Figure 2A and Figure S2A). In *Arabidopsis*, seven LOG enzymes with phosphoribohydrolase activity can directly convert iPRMP into iP (Kuroha et al. 2009; Tokunaga et al., 2011). Thus, elevated levels of iPRPs in the shoot likely serve as a reservoir of latent CK activity, with the potential to regulate growth and gene expression depending on the spatial and temporal activity of LOG enzymes. Given that Arabidopsis *LOG* genes exhibit distinct spatial expression patterns (Kuroha et al. 2009), the iPRPs accumulated in response to shoot nitrate accumulation may be locally converted to iP in cells or tissues expressing specific LOG isoforms. While our CK quantifications were performed using whole shoots and roots, it is possible that localized changes in iP concentrations —for example, in actively growing regions such as the shoot apical meristem or the basal zone of young leaves—were not captured. Targeted sampling of these regions may therefore be necessary to reveal the spatial dynamics of iP production in response to shoot nitrate status.

**Figure 2.**
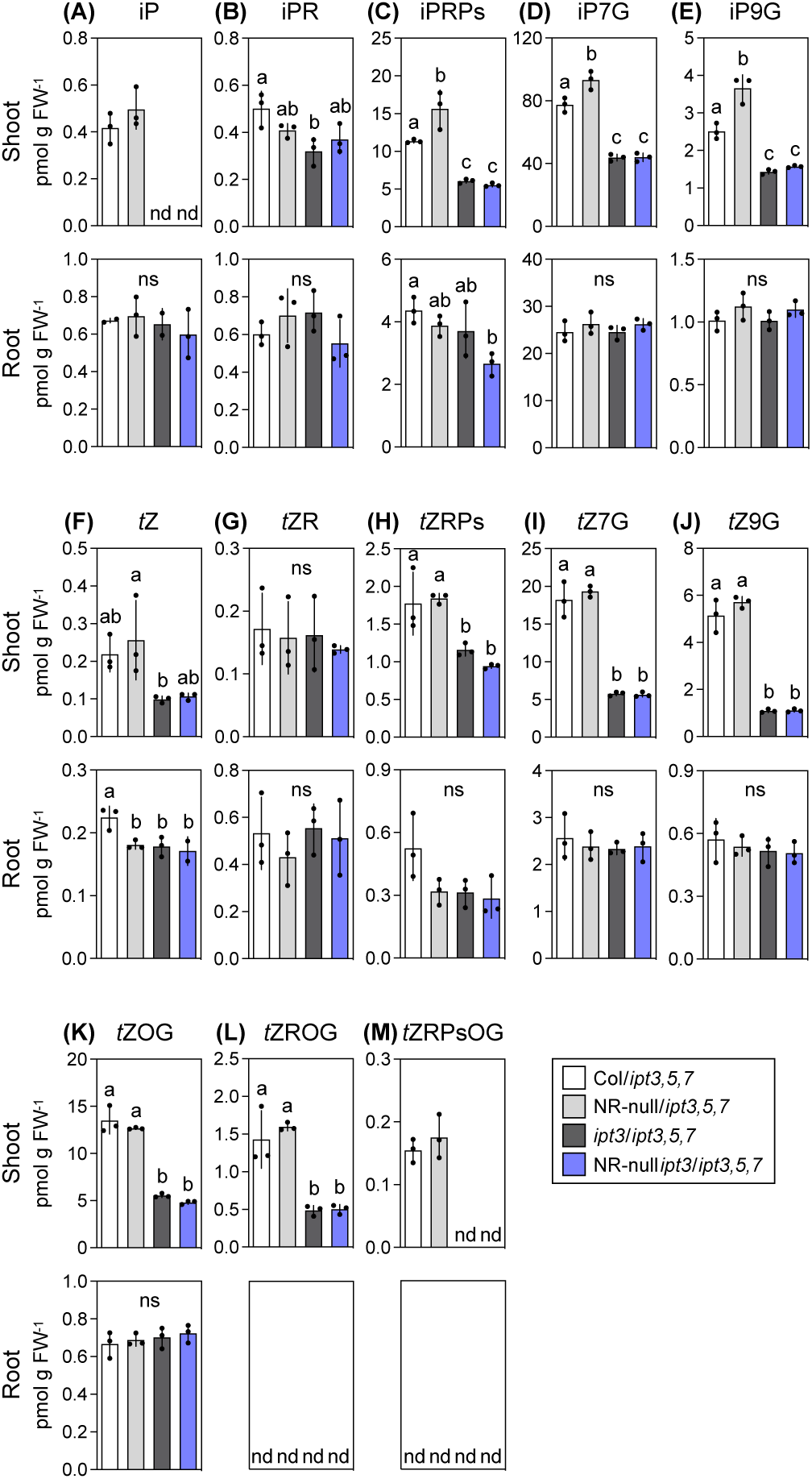
Effects of shoot nitrate status and shoot *IPT3* expression on the shoot and root concentrations of CK species. (A–M) The shoot and root concentrations of iP (A), iPR (B), iPRPs (C), iP7G (D), iP9G (E), *t*Z (F), *t*ZR (G), *t*ZPRs (H), *t*Z7G (I), *t*Z9G (J), *t*ZOG (K), *t*ZROG (L), and *t*ZRPsOG (M) in the grafted plants 5 days after nitrogen removal (condition 5). Sampling was performed three times independently. In the sampling, 7, 6, and 5 grafted plants were pooled as one biological replicate. Data are presented as mean ± SD [n = 3, excluding iP concentrations in Col/*ipt3,5,7* roots and *ipt3*/*ipt3,5,7* roots (n = 2)]. Different lowercase letters indicate significant differences, as determined via Tukey–Kramer test (*P* < 0.05). iP, *N*^6^-(Δ^2^-isopentenyl)adenine; iPR, iP riboside; iPRPs, iP ribotides; iP7G, iP-7-*N*-glucoside; iP9G, iP-9-*N*-glucoside; *t*Z, *trans*-zeatin; *t*ZR, *t*Z riboside; *t*ZRPs, *t*Z ribotides; *t*Z7G, *t*Z-7-*N*-glucoside; *t*Z9G, *t*Z-9-*N*-glucoside; *t*ZOG, *t*Z-*O*-glucoside; *t*ZROG, *t*ZR-*O*-glucoside; *t*ZRPsOG, *t*ZRPs-*O*-glucoside; ns, not significant; nd, not detected; FW, fresh weight.

### Shoot nitrate accumulation enhances shoot growth via shoot IPT3

To determine the effects of shoot nitrate status and shoot *IPT3* expression on plant growth, we analyzed growth parameters using all grafted plants from five independent experiments (Figure 3). Shoot fresh weight, leaf blade area, rosette diameter, and leaf number were significantly lower in *ipt3*/*ipt3,5,7* and NR-null *ipt3*/*ipt3,5,7* plants than in Col/*ipt3,5,7* and NR-null/*ipt3,5,7* plants (Figure 3B–E). Meanwhile, no significant difference was observed in the root fresh weight among the grafted plants (Figure 3F). These observations indicate that shoot IPT3 generally contributes to shoot growth. Importantly, shoot appearance, shoot fresh weight, and leaf blade area were significantly increased by shoot NR deficiency in the presence of shoot IPT3 but not in its absence (Figure 3A–C). A similar trend was observed for the rosette diameter, although the differences were not statistically significant (Figure 3D). These patterns correspond to the findings for shoot iPRPs, iP7G, and iP9G concentrations (Figure 2C–E and Figure S2C–E). Actually, shoot fresh weight and leaf blade area were strongly correlated with levels of these iP-type CKs (Figure S3). In *Arabidopsis*, iPRPs and the glucosides iP7G and iP9G are generally considered biologically inactive CKs (Tokunaga et al. 2011; Hoyerová and Hošek 2020) and thus unlikely to directly promote shoot growth. Among iPRPs, iPRMP can be rapidly converted into bioactive iP by LOG enzymes, thereby activating CK signaling (Kurakawa et al. 2007; Kuroha et al. 2009). By contrast, iP7G and iP9G are not converted into other CKs (Hošek et al. 2020), precluding their contribution to shoot growth. Nevertheless, because bioactive iP is predominantly metabolized to iP7G (Hošek et al., 2020) and iP7G resists degradation by Arabidopsis CKXs (Galuszka et al., 2007), iP7G may serve as a proxy for iP-dependent signaling. Overall, our findings suggest that shoot nitrate accumulation promotes shoot growth by increasing IPT3 and iP-type CK biosynthesis in the shoot. It is well established that in Arabidopsis plants, root-supplied nitrate stimulates the biosynthesis of *t*Z-type CKs in the root and facilitates their transport to the shoot, thereby promoting shoot growth (Kiba et al. 2013; Ko et al. 2014; Osugi et al. 2017; Sakakibara, 2021). Collectively, shoot-derived iP-type CKs and root-derived *t*Z-type CKs may regulate shoot growth by responding to the internal and external nitrate status, respectively. Interestingly, the fresh weight ratios of shoot to root were significantly elevated by shoot NR deficiency in the presence of shoot IPT3 but not in its absence (Figure 3G). Given that, in general, biomass ratios of shoot to root are positively correlated with overall nitrogen availability both inside and outside the plant (Andrews et al. 2006; Krapp et al. 2011; Hachiya et al. 2014), our findings support the idea that shoot nitrate status serves as an indicator of nitrogen availability. Furthermore, since CK action is known to enhance sink activity (Werner et al. 2008), it is plausible that shoot nitrate status influences shoot CK levels and subsequently biomass allocation via shoot IPT3-mediated CK signaling.

**Figure 3.**
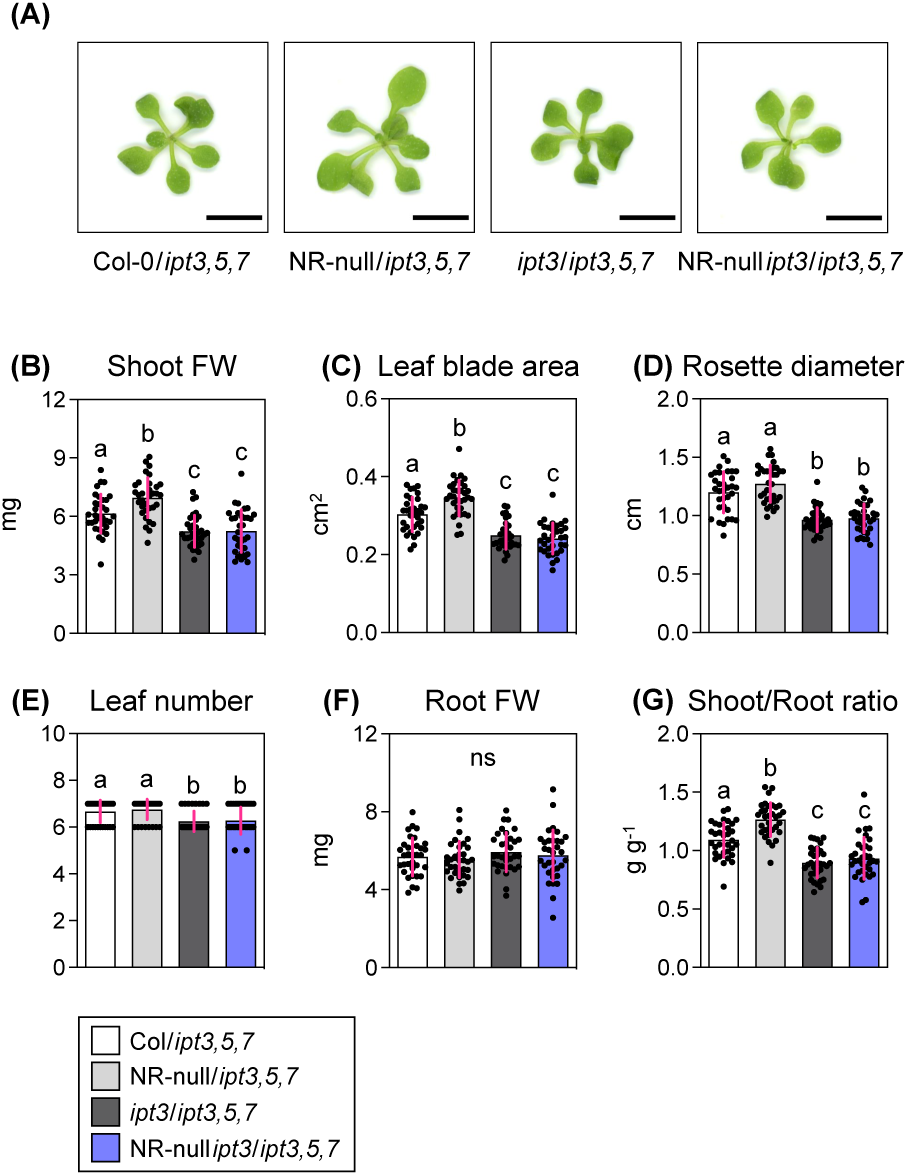
Effects of shoot nitrate status and shoot *IPT3* expression on plant growth. (A) Representative shoot appearance of grafted plants. Scale bars represent 5 mm. (B–F) Shoot fresh weight (B), leaf blade area (C), rosette diameter (D), leaf number (E), root fresh weights (F), and fresh weight ratios of shoot to root (G) of grafted plants 5 days after nitrogen removal (condition 5). All data pooled from five independent grafting experiments are presented as mean ± SD (n = 33). Different lowercase letters indicate significant differences, as determined via Tukey–Kramer test (*P* < 0.05). ns, not significant; FW, fresh weight.

### Shoot IPT3 oppositely modulates transcriptomic responses to shoot nitrate accumulation in shoots and roots

To investigate the effects of shoot nitrate status and shoot *IPT3* expression on systemic responses, we performed RNA-seq using the shoots and roots of grafted plants, with three biological replicates collected independently. Reads per million mapped reads (RPM; Table S2) and normalized transcript levels (Table S3) were calculated according to the read counts. To identify genes regulated by shoot nitrate accumulation and further modulated by shoot IPT3, we extracted differentially expressed genes (DEGs) between NR-null/*ipt3,5,7* and Col/*ipt3,5,7* and between NR-null *ipt3*/*ipt3,5,7* and *ipt3*/*ipt3,5,7* with a minimum fold change of 1.5 (Figure 4A and Figure S4 and Table S4). Comparison of these DEGs between shoots and roots revealed minimal overlap (Figure S4), indicating that shoots and roots express distinct sets of genes regulated by shoot nitrate accumulation. Hence, we analyzed the shoot and root transcriptomes separately. Notably, in shoots, the number of DEGs was larger between NR-null/*ipt3,5,7* and Col/*ipt3,5,7* than between NR-null *ipt3*/*ipt3,5,7* and *ipt3*/*ipt3,5,7* (Figure 4A). By contrast, in roots, the number of DEGs was lower between NR-null/*ipt3,5,7* and Col/*ipt3,5,7* than between NR-null *ipt3*/*ipt3,5,7* and *ipt3*/*ipt3,5,7* (Figure 4A). These observations suggest that shoot transcriptional responses to shoot nitrate accumulation are enhanced by shoot IPT3, whereas root responses are attenuated by shoot IPT3. To substantiate this, we quantitatively analyzed the 631 and 663 DEGs identified exclusively between NR-null/*ipt3,5,7* and Col/*ipt3,5,7* in shoots, and the 189 and 126 DEGs identified exclusively between NR-null *ipt3*/*ipt3,5,7* and *ipt3*/*ipt3,5,7* in roots (Figure 4A). The shoot transcriptional responses of 631 upregulated and 663 downregulated DEGs to shoot NR deficiency were decreased by shoot IPT3 deficiency (Figure 4B, C). Conversely, the root responses of 189 upregulated and 126 downregulated DEGs were augmented by shoot IPT3 deficiency (Figure 4D, E). Furthermore, k-means clustering identified 11 out of 16 clusters (clusters 1–3, 6–8, 11, 12, and 14–16) in which gene expression was significantly altered by shoot NR deficiency, with their responses decreased by shoot IPT3 deficiency in shoots but increased in roots (Figure S5, S6 and Table S5). In summary, we conclude that shoot transcriptional responses to shoot nitrate accumulation are generally enhanced by shoot IPT3, whereas root responses are dampened by shoot IPT3. It should be noted that three independent RNA-seq results exhibited similar trends; therefore, only the mean transcript levels are presented in subsequent figures for clarity, unless otherwise stated.

**Figure 4.**
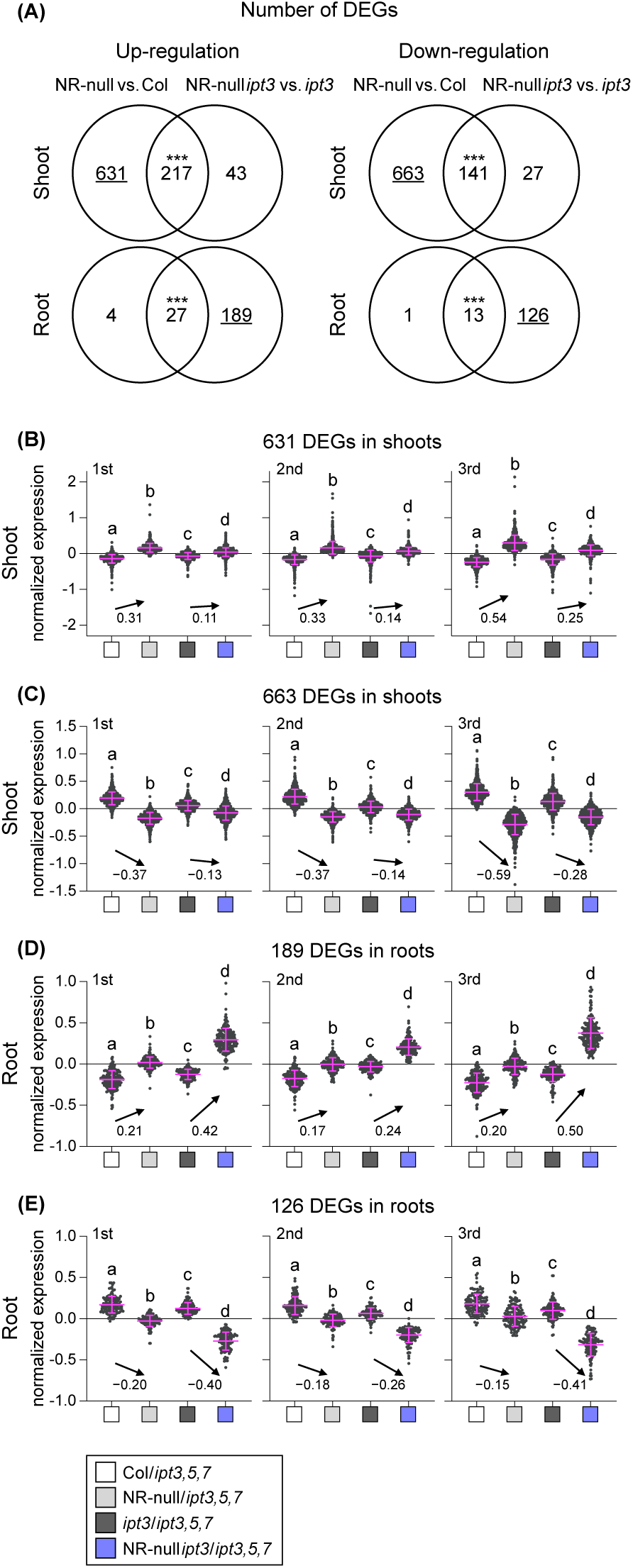
Effects of shoot nitrate status and shoot *IPT3* expression on the shoot and root transcriptomes. RNA-seq was performed using the shoots and roots of grafted plants 5 days after nitrogen removal (condition 5). Sampling was performed three times independently. In the sampling, 9, 6, and 7 grafted plants were pooled as one biological replicate. (A) Venn diagram presenting the number of DEGs at a minimum fold change of 1.5 in NR-null/*ipt3,5,7* plants relative to Col/*ipt3,5,7* plants and in NR-null *ipt3*/*ipt3,5,7* plants relative to *ipt3*/*ipt3,5,7* plants. Statistical significance of overlap was assessed using the hypergeometric test (\**P* < 0.05; \*\**P* < 0.01; \*\*\**P* < 0.001). (B–E) Plots of normalized transcript levels of 631 DEGs (B) and 663 DEGs (C) in shoots and 189 DEGs (D) and 126 DEGs (E) in roots. 1st, 2nd, and 3rd denote the order of three independent experiments. Different lowercase letters indicate significant differences, as determined via Tukey–Kramer test (*P* < 0.05). The numbers on the graph represent the difference in mean values.

Enrichment analyses of the 11 clusters from Figure S5 and S6 revealed a diverse set of genes significantly regulated by shoot nitrate and shoot IPT3. In shoots, the cluster 6 and 8 enriched terms related to translation, primary carbon metabolism, and biotic stress response such as “ath03010: Ribosome”, “WP2621: Glycolysis”, and “GO:0009617: response to bacterium” (Figure S5A). Previously, we reported that shoot nitrate accumulation is accompanied by the upregulation of ribosome-associated genes in shoots (Hachiya et al. 2020). It is widely known that CK signal upregulates genes encoding ribosomal proteins and elongation factors (Sakakibara et al. 2006). Hence, translation may be activated by shoot nitrate accumulation, some of which are mediated by shoot IPT3 (Figure S5B). Meanwhile, terms related to wounding and jasmonic acids were overrepresented in both cluster 3 and 7 (Figure S5A). Thus, stress responses associated with jasmonic acids signaling could be downregulated by shoot nitrate accumulation, which is enhanced by shoot IPT3 (Figure S5B). In roots, the cluster 11, 12, and 14 predominantly featured terms related to abiotic and biotic stresses, particularly hypoxia (Figure S6A). The cluster 15 and 16 enriched terms associated with translation, biosynthesis, and nitrate transport (Figure S6A). Notably, the genes encoding high-affinity nitrate transporters (*NRT2.1*, *NRT2.4*, *NRT2.5*, *NRT3.1*) and nitrate reductases (*NIA1*, *NIA2*) were included in the cluster 16 (Table S5). Thus, shoot nitrate accumulation may suppress nitrate uptake and assimilation in the roots, the responses of which are dampened by shoot IPT3 (Figure S6B).

### Shoot nitrate accumulation induces shoot *ARR2* expression via shoot IPT3

To investigate CK responses associated with altered shoot iP-type CKs and shoot growth, we first examined the shoot expression of representative CK-inducible genes using a reference list compiled from multiple studies (Bhargava et al. 2013). However, their expression showed statistically significant but slight decreases due to shoot IPT3 deficiency, whereas no significant changes were detected with shoot NR deficiency (Figure S7A and Table S6). Notably, previous studies analyzed early transcriptomic responses to exogenous application of *t*Z and benzyladenine (Bhargava et al. 2013); therefore, conventional gene lists might be unsuitable for evaluating steady-state responses to alterations of endogenous iP-type CKs. Next, to further dissect CK-related components, we focused on CK signaling- and metabolism-related genes, including those encoding ARRs, cytokinin response factors (CRFs), cytochrome P450 monooxygenases (CYP735A1/A2), UGTs, and CKXs (Werner et al. 2003; Rashotte and Goertzen 2010; Kiba et al. 2013; Hošek et al. 2020) (Figure S7B). Among these, shoot expression of the type-B *ARR2* was induced by shoot NR deficiency in the presence of shoot IPT3 but not in its absence (Figure S7B), a finding corroborated by RT-qPCR (Figure S7C). Choi et al. (2010) demonstrated that CK-activated ARR2 enhances Arabidopsis immunity through direct binding with the transcription factor TGA3. In Figure S5A, the cluster 8 derived from clustering was overrepresented in terms such as “GO:0009617: response to bacterium” and “GO:0009627: systemic acquired resistance” (Figure S5A). Thus, shoot nitrate accumulation might enhance immunity by upregulating *ARR2* via shoot IPT3. Furthermore, given that *ARR2*-overexpressing plants exhibit larger rosette leaves and biomass (Arnaud et al. 2017), it is plausible that shoot nitrate accumulation could promote shoot growth partly via shoot IPT3–ARR2 module.

### Shoot IPT3 enhances transcriptional responses of nitrate–CPKs–NLPs-regulatory genes to shoot nitrate accumulation

Previously, we reported that the expression of nitrate-responsive genes is influenced by shoot nitrate accumulation (Okamoto et al. 2019), prompting our focus on these genes. In shoots, the transcript levels of genes induced by nitrate itself (Wang et al. 2004) were significantly increased by shoot NR deficiency, and this induction was attenuated by shoot IPT3 deficiency (Figure 5A and Table S7). By contrast, there was minimal variation in the root expression of nitrate-inducible genes among the plants. Given that CPK10/30/32, NLP7, and its homologs (NLPs) govern nitrate-dependent gene expression (Liu et al. 2017; Liu et al. 2022), we analyzed the expression of genes induced by the nitrate–CPKs–NLPs signaling pathway. In shoots, the genes induced via CPK10/30/32 and those co-induced via all NLPs exhibited similar expression patterns as the nitrate-inducible genes (Figure 5A–C and Table S7). Meanwhile, none of the genes induced exclusively by individual NLPs exhibited different expression patterns among the plants (Figure S8 and Table S8). In roots, the expression of genes regulated via CPK10/30/32 and NLPs showed little variation among the plants (Figure 5B, C and Figure S8 and Table S7, S8). These findings suggest that shoot nitrate accumulation upregulates nitrate-inducible genes via CPK10/30/32 and multiple NLPs specifically in the shoots, with their responses enhanced by shoot IPT3.

**Figure 5.**
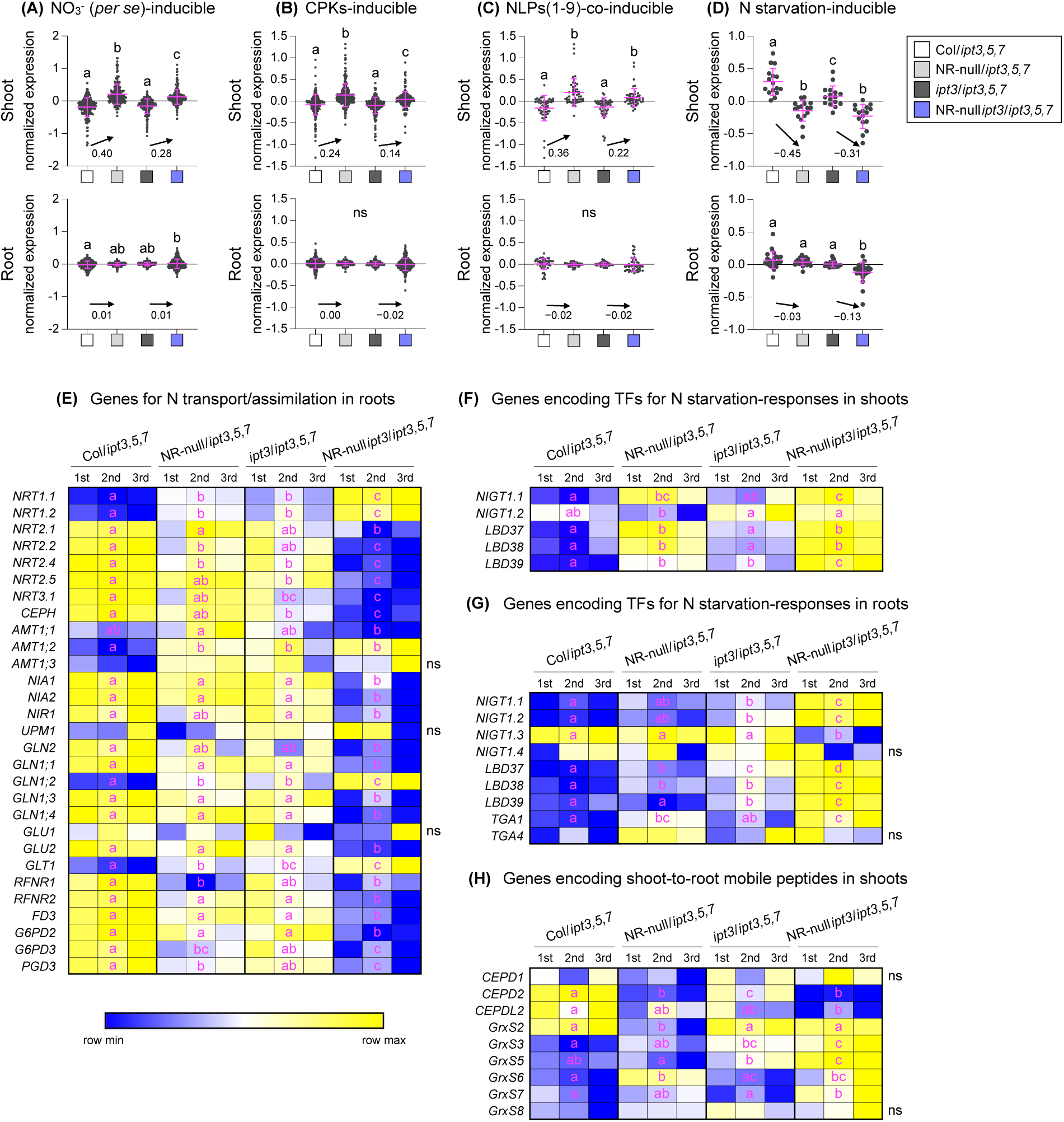
Effects of shoot nitrate status and shoot *IPT3* expression on nitrogen-related transcriptional responses in shoots and roots. (A–D) Plots of mean normalized transcript levels of nitrate (*per se*)-inducible genes in 2 h (A), genes induced via CPK10/30/32 (B), genes commonly induced via all NLPs (C), and nitrogen starvation-inducible genes (D) in the shoots and roots of grafted plants 5 days after nitrogen removal (condition 5). The gene lists were obtained from Wang et al. (2004) for (A), Liu et al. (2017) for (B), Liu et al. (2022) for (C), and Kiba et al. (2018) and Krapp et al. (2011) for (D). The numbers on the graph represent the difference in mean values. (E–H) Transcriptional changes in genes involved in nitrogen transport and assimilation in roots (E), genes encoding transcription factors for nitrogen starvation responses in shoots (F) and in roots (G), and genes encoding shoot-to-root mobile peptides in shoots (H) in grafted plants 5 days after nitrogen removal. The normalized transcript levels were visualized as heatmaps using MORPHEUS (https://software.broadinstitute.org/morpheus). The relative color scheme uses the minimum (blue) and maximum (yellow) values in each row. 1st, 2nd, and 3rd denote the order of three independent experiments. TF, transcription factor. (A–H) Different lowercase letters indicate significant differences, as determined via Tukey–Kramer test (*P* < 0.05). ns, not significant.

### Shoot IPT3 oppositely modulates transcriptional responses of nitrogen starvation-inducible genes to shoot nitrate accumulation in shoots and roots

Previously, we reported that shoot nitrate accumulation alleviates nitrogen starvation responses (Okamoto et al. 2019). Thus, we analyzed the responses of two independent sets of nitrogen starvation-inducible gene (Krapp et al. 2011; Kiba et al. 2018) (Table S9). Although these gene sets are quite distinct (Figure S9A), shoot NR deficiency consistently repressed shoot expression of nitrogen starvation-inducible genes, with this repression slightly alleviated by shoot IPT3 deficiency (Figure S9B, C). The nitrogen starvation-inducible genes common to both gene sets exhibited a similar but stronger tendency (Figure 5D). Meanwhile, the root expression of nitrogen starvation inducible-genes were significantly repressed by shoot NR deficiency only in the absence of shoot IPT3 (Figure 5D and S9B, C). Overall, upon nitrate accumulation in shoots, shoot IPT3 enhances the repression of nitrogen starvation-inducible genes in shoots but attenuates this repression in roots.

Several genes involved in nitrogen transport and assimilation in the root are often induced under nitrogen starvation (Krapp et al. 2011; Kiba et al. 2018), and thus, we focused on their root expression. Most of the genes involved in high-affinity nitrate uptake and nitrogen assimilation were slightly downregulated by shoot NR deficiency, and these responses were remarkably intensified by shoot IPT3 deficiency (Figure 5E). Thus, the degree of downregulation in root nitrate utilization due to shoot nitrate accumulation is largely dependent on shoot IPT3. By contrast, genes involved in dual-affinity/low-affinity nitrate transport (*NRT1.1* and *NRT1.2*) (Liu et al. 1999) were upregulated by shoot NR deficiency regardless of the presence of shoot IPT3 (Figure 5E). Because NRT1.1 and NRT1.2 can facilitate the uptake of auxin and abscisic acid, respectively (Krouk et al. 2010; Kanno et al. 2012), their upregulation may not necessarily be related to nitrate acquisition.

### Systemic regulation of nitrogen starvation-inducible genes by shoot nitrate accumulation may involve transcriptional repressors and mobile peptides

Nitrate signaling induces the expression of genes encoding Nitrate-inducible GARP-type transcriptional repressor 1 family proteins (NIGT1.1/1.2/1.3/1.4) and LOB domain-containing proteins 37/38/39 (LBD37/38/39) via NLPs (Rubin et al. 2009; Marchive et al. 2013; Kiba et al. 2018; Maeda et al. 2018; Safi et al 2021). These proteins act as repressors of nitrogen starvation-inducible genes; therefore, we examined their encoding genes. In the shoot, the transcript levels of *NIGT1.1* and *LBD37/38* were increased by shoot NR deficiency, and this induction was attenuated by shoot IPT3 deficiency (Figure 5F and S10A). The expression of these repressors exhibited a strong negative correlation with that of nitrogen starvation-inducible genes in the shoot (Figure S10B). In the root, the transcript levels of *NIGT1.1/1.2* and *LBD37/38* were increased by shoot NR deficiency, and this induction was enhanced by shoot IPT3 deficiency (Figure 5G and S10C). Interestingly, root *LBD39* expression was induced by shoot NR deficiency only in the absence of shoot IPT3. These repressors also showed a strong negative correlation with nitrogen starvation-inducible genes in the root (Figure S10D). These findings suggest that shoot nitrate accumulation and shoot IPT3 regulate the systemic regulation of nitrogen starvation-inducible genes via these repressors. It is valuable to explore whether iP-type CKs synthesized by shoot IPT3 systemically regulate the root expression of repressor genes.

Next, we focused on candidate shoot-to-root mobile peptides regulating systemic nitrogen responses, i.e., CEPD1/2, CEPDL2, and GrxS1–8 (Ohkubo et al. 2017; Ota et al. 2020; Kobayashi et al. 2024) (Figure 5H and S11). The co-activator complexes composed of CEPD1/2/CEPDL2 and TGA1/4 promote root nitrate uptake via upregulation of *NRT2.1* and *CEPH*, whereas the co-repressor complexes of GrxS1–8 and TGA1/4 repress it (Kobayashi et al. 2024). In the shoot, *CEPDL2* transcript levels were reduced by shoot NR deficiency, and this repression was attenuated by shoot IPT3 deficiency (Figure 5H and S11A). In contrast, shoot *GrxS3* expression was upregulated by shoot NR deficiency, and this induction was slightly enhanced in the absence of shoot IPT3. Interestingly, shoot *GrxS5* expression was induced by shoot NR deficiency only in the absence of shoot IPT3. The root expression of *NRT2.1* and *CEPH* was positively correlated with shoot expression of *CEPDL2*, whereas negatively correlated with shoot expression of *GrxS3* and *GrxS5* (Figure S11B). These findings suggest that shoot nitrate status and shoot IPT3 regulate root high-affinity nitrate uptake via shoot-to-root mobile peptides.

### Conclusion: a scheme integrating shoot nitrate accumulation responses with shoot IPT3

Our experimental system elucidated the systemic responses to shoot nitrate status and their dependence on shoot IPT3. A novel scheme for the systemic responses is summarized in Figure 6: shoot nitrate accumulation promotes shoot growth and induces the shoot expression of *IPT3*, nitrate–CPKs–NLPs-inducible genes (including those for nitrate uptake/assimilation), and immune response genes, with their responses enhanced by shoot IPT3. Meanwhile, shoot nitrate accumulation decreases the root expression of nitrogen starvation-inducible genes and nitrate uptake/assimilation genes, with their responses dampened by shoot IPT3. These interactive effects of shoot nitrate status and shoot IPT3 were confirmed by two-way ANOVA, which detected statistically significant interactions between the presence or absence of shoot NR and shoot IPT3 (Table S10). Hence, under shoot nitrate accumulation, shoot IPT3 would promote nitrogen utilization at the whole-plant scale, providing organic nitrogen crucial for shoot growth enhancement. Notably, plant CK levels are generally reduced by prolonged abiotic stresses including salinity, drought, chilling, and heat (Cortleven et al. 2019), during which shoot growth is usually suppressed. Thus, shoot growth and nitrogen utilization, which are tuned by shoot IPT3 depending on shoot nitrate status, would be further fine-tuned via modulation of CK levels depending on abiotic stresses. Such crosstalk in CK signaling would optimize plant growth and nitrogen utilization in fluctuating environments. Despite these insights, it remains challenging to distinguish between primary and secondary responses to shoot nitrate accumulation, as well as to evaluate the potential effects of organic nitrogen compounds such as Gln, a candidate nitrogen status signal (Nacry et al., 2013), which were not quantified in this study. In addition, it is unclear exactly where and how the phloem-mobile iP-type CKs produced by shoot IPT3 trigger downstream signaling to orchestrate systemic responses. Furthermore, the cell types, subcellular compartments, and molecular mechanisms involved in sensing shoot nitrate status are entirely unknown. Future studies should address these unresolved questions.

**Figure 6.**
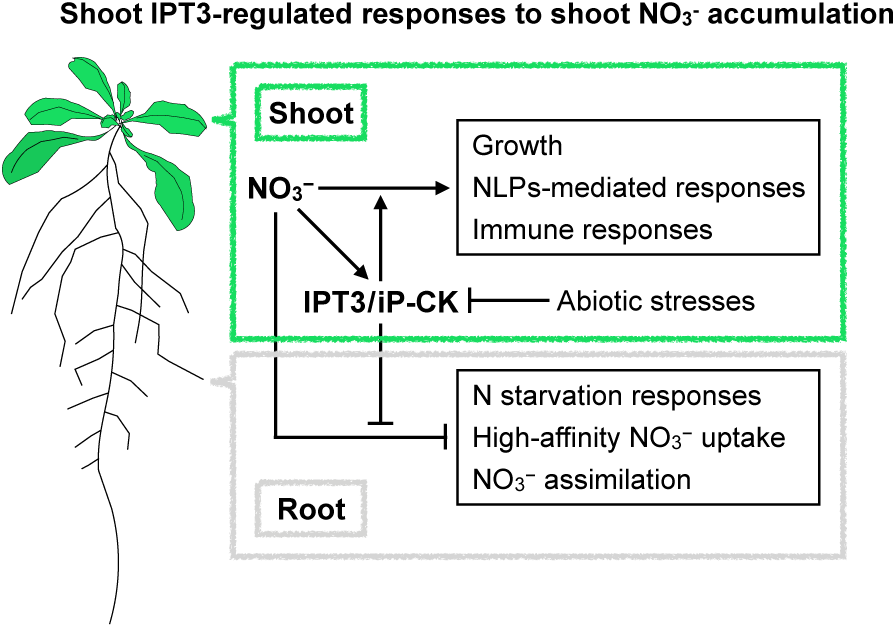
Summary of systemic responses to shoot nitrate accumulation and their dependence by shoot IPT3.

## Author contributions

TH: conceptualization; TS: data curation; KM and TH: formal analysis; KM and TH: funding acquisition; KM, TS, MK, YT, and DS: investigation; KM: methodology; KM and TH: project administration; KM and TH: resources; TN, HS, and TH: supervision; KM and TH: validation; KM and TH: visualization; TH: writing – original draft preparation; All authors: writing – review & editing.

## Acknowledgements

The NR-null seeds were kindly provided by Dr. Nigel M. Crawford (University of California, San Diego). The *ipt3* and *ipt3,5,7* seeds were kindly provided by Dr. Tatsuo Kakimoto (Osaka University). We thank Ms. Yuki Okamoto and Mr. Masahiro Watanabe for their support in the preparation of the plant materials. This study was supported by JSPS KAKENHI [No. 20K05771 and 23K04978 to TH and No. JP24KJ1709 to KM], by JST SPRING [No. JPMJSP2155 to KM], and by the Sasakawa Scientific Research Grant from the Japan Science Society [No. 2022-4039 to KM].

## Data Availability Statement

The RNA-seq raw data are available in ArrayExpress under the accession number E-MTAB-14302 (https://www.ebi.ac.uk/biostudies/arrayexpress).

## Supporting Information

Additional supporting information can be found online in the Supporting Information section.

## References

Alvarez, J. M., E. Riveras, E. A. Vidal, et al. 2014. “Systems approach identifies TGA1 and TGA4 transcription factors as important regulatory components of the nitrate response of *Arabidopsis thaliana* roots.” Plant Journal 80 : 1–13. 10.1111/tpj.12618.

Andrews, M., J. A. Raven, P. J. Lea, and J. I. Sprent. 2006. “A role for shoot protein in shoot–root dry matter allocation in higher plants.” Annals of Botany 97: 3–10. 10.1093/aob/mcj009.

Arnaud, D., S. Lee, Y. Takebayashi, et al. 2017. “Cytokinin-mediated regulation of reactive oxygen species homeostasis modulates stomatal immunity in Arabidopsis.” Plant Cell 29: 543–559. 10.1105/tpc.16.00583.

Bhargava, A., I. Clabaugh, J. P. To, et al. 2013. “Identification of cytokinin-responsive genes using microarray meta-analysis and RNA-Seq in Arabidopsis.” Plant Physiology 162: 272–294. 10.1104/pp.113.217026.

Bishopp, A., S. Lehesranta, A. Vatén, et al. 2011. “Phloem-transported cytokinin regulates polar auxin transport and maintains vascular pattern in the root meristem.” Current Biology 21: 927–932. 10.1016/j.cub.2011.04.049.

Bouguyon, E., F. Brun, D. Meynard, et al. 2015. “Multiple mechanisms of nitrate sensing by *Arabidopsis* nitrate transceptor NRT1.1.” Nature Plants 1: 15015. 10.1038/nplants.2015.15.

Choi, J., S. U. Huh, M. Kojima, H. Sakakibara, K. H. Paek, and I. Hwang. 2010. “The cytokinin-activated transcription factor ARR2 promotes plant immunity via TGA3/NPR1-dependent salicylic acid signaling in *Arabidopsis*.” Developmental Cell 19: 284–295. 10.1016/j.devcel.2010.07.011.

Cortleven, A., J. E. Leuendorf, M. Frank, D. Pezzetta, S. Bolt, and T. Schmülling. 2019. “Cytokinin action in response to abiotic and biotic stresses in plants.” Plant, Cell & Environment 42: 998–1018. 10.1111/pce.13494.

Durand, M., V. Brehaut, G. Clement, et al. 2023. “The Arabidopsis transcription factor NLP2 regulates early nitrate responses and integrates nitrate assimilation with energy and carbon skeleton supply.” Plant Cell 35: 1429–1454. 10.1093/plcell/koad025.

Galuszka P., H. Popelková, T. Werner, et al. 2007. “Biochemical characterization of cytokinin oxidases/dehydrogenases from *Arabidopsis thaliana* expressed in *Nicotiana tabacum* L.” Journal of Plant Growth Regulation 26: 255–267.

Ge, S. X., E. W. Son, and R. Yao. 2018. “iDEP: an integrated web application for differential expression and pathway analysis of RNA-Seq data.” BMC Bioinformatics 19: 534. 10.1186/s12859-018-2486-6.

Hachiya, T., D. Sugiura, M. Kojima, et al. 2014. “High CO_2_ triggers preferential root growth of *Arabidopsis thaliana* via two distinct systems under low pH and low N stresses.” Plant & Cell Physiology 55: 269–280. 10.1093/pcp/pcu001.

Hachiya, T., N. Ueda, M. Kitagawa, et al. 2016. “Arabidopsis root-type ferredoxin:NADP(H) oxidoreductase 2 is involved in detoxification of nitrite in roots.” Plant & Cell Physiology 57: 2440–2450. 10.1093/pcp/pcw158.

Hachiya, T., and H. Sakakibara. 2017. “Interactions between nitrate and ammonium in their uptake, allocation, assimilation, and signaling in plants.” Journal of Experimental Botany 68: 2501–2512. 10.1093/jxb/erw449.

Hachiya, T., Y. Okamoto, M. Watanabe, et al. 2020. “Genome-wide responses to shoot nitrate satiety are attenuated by external ammonium in *Arabidopsis thaliana*.” Soil Science and Plant Nutrition 66: 317–327. 10.1080/00380768.2020.1717905.

Hachiya, T., T. Oya, K. Monden, A. Nagae, and T. Nakagawa. 2021a. “A cellophane-supported *Arabidopsis* culture for seamless transfer between different media is useful for studying various nitrogen responses.” Soil Science and Plant Nutrition 67: 277–282. 10.1080/00380768.2021.1908094.

Hachiya, T., J. Inaba, M. Wakazaki, et al. 2021b. “Excessive ammonium assimilation by plastidic glutamine synthetase causes ammonium toxicity in *Arabidopsis thaliana*.” Nature Communications 12: 4944. 10.1038/s41467-021-25238-7.

Hirose, N., K. Takei, T. Kuroha, T. Kamada-Nobusada, H. Hayashi, and H. Sakakibara. 2008. “Regulation of cytokinin biosynthesis, compartmentalization and translocation.” Journal of Experimental Botany 59: 75–83. 10.1093/jxb/erm157.

Ho, C. H., S. H. Lin, H. C. Hu, and Y. F. Tsay. 2009. “CHL1 functions as a nitrate sensor in plants.” Cell 138: 1184–1194. 10.1016/j.cell.2009.07.004.

Hošek P., K. Hoyerová, N. S. Kiran, et al. 2020. “Distinct metabolism of *N*-glucosides of isopentenyladenine and *trans*-zeatin determines cytokinin metabolic spectrum in Arabidopsis.” New Phytologist 225: 2423–2438. 10.1111/nph.16310.

Hoyerová, K., P. Hošek. 2020. “New insights into the metabolism and role of cytokinin *N*-glucosides in plants.” Frontiers in Plant Science 11: 741. 10.3389/fpls.2020.00741.

Hu, H. C., Y. Y. Wang, and Y. F. Tsay. 2009. “AtCIPK8, a CBL-interacting protein kinase, regulates the low-affinity phase of the primary nitrate response.” Plant Journal 57: 264–278. 10.1111/j.1365-313X.2008.03685.x.

Iwagami, T., T. Ogawa, T. Ishikawa, and T. Maruta. 2022. “Activation of ascorbate metabolism by nitrogen starvation and its physiological impacts in *Arabidopsis thaliana*.” Bioscience, Biotechnology, and Biochemistry 86: 476–489. 10.1093/bbb/zbac010.

Kanno, Y., A. Hanada, Y. Chiba, et al. 2012. “Identification of an abscisic acid transporter by functional screening using the receptor complex as a sensor.” Proceedings of the National Academy of Sciences of the United States of America 109: 9653–9658. 10.1073/pnas.1203567109.

Kiba, T., K. Takei, M. Kojima, and H. Sakakibara. 2013. “Side-chain modification of cytokinins controls shoot growth in *Arabidopsis*.” Developmental Cell 27: 452–461. 10.1016/j.devcel.2013.10.004.

Kiba, T., J. Inaba, T. Kudo, et al. 2018. “Repression of nitrogen starvation responses by members of the Arabidopsis GARP-type transcription factor NIGT1/HRS1 subfamily.” Plant Cell 30: 925–945. 10.1105/tpc.17.00810.

Ko, D., J. Kang, T. Kiba, et al. 2014. “*Arabidopsis* ABCG14 is essential for the root-to-shoot translocation of cytokinin.” Proceedings of the National Academy of Sciences of the United States of America 111: 7150–7155. 10.1073/pnas.1321519111.

Kobayashi, R., Y. Ohkubo, M. Izumi, et al. 2024. “Integration of shoot-derived polypeptide signals by root TGA transcription factors is essential for survival under fluctuating nitrogen environments.” Nature Communications 15: 6903. 10.1038/s41467-024-51091-5.

Kojima, M., T. Kamada-Nobusada, H. Komatsu, et al. 2009. “Highly sensitive and high-throughput analysis of plant hormones using MS-probe modification and liquid chromatography–tandem mass spectrometry: an application for hormone profiling in *Oryza sativa*.” Plant & Cell Physiology 50: 1201–1214. 10.1093/pcp/pcp057.

Kojima, M., N. Makita, K. Miyata, et al. 2023. “A cell wall–localized cytokinin/purine riboside nucleosidase is involved in apoplastic cytokinin metabolism in *Oryza sativa*.” Proceedings of the National Academy of Sciences of the United States of America 120: e2217708120. 10.1073/pnas.221770812.

Konishi, M., and S. Yanagisawa. 2013. “Arabidopsis NIN-like transcription factors have a central role in nitrate signalling.” Nature communications 4: 1617. 10.1038/ncomms2621.

Krapp, A., R. Berthomé, M. Orsel, et al. 2011. “Arabidopsis roots and shoots show distinct temporal adaptation patterns toward nitrogen starvation.” Plant Physiology 157: 1255–1282. 10.1104/pp.111.179838.

Krouk, G, B. Lacombe, A. Bielach, et al. 2010. “Nitrate-regulated auxin transport by NRT1.1 defines a mechanism for nutrient sensing in plants.” Developmental Cell 18: 927–937. 10.1016/j.devcel.2010.05.008.

Kurakawa T., N. Ueda, M. Maekawa, et al. 2007. “Direct control of shoot meristem activity by a cytokinin-activating enzyme.” Nature 445: 652–655. 10.1038/nature05504.

Kuroha, T., H. Tokunaga, M. Kojima, et al. 2009. “Functional analyses of *LONELY GUY* cytokinin-activating enzymes reveal the importance of the direct activation pathway in *Arabidopsis*.” Plant Cell 21: 3152–3169. 10.1105/tpc.109.068676.

Langmead, B., C. Trapnell, M. Pop, and S. L. Salzberg. 2009. “Ultrafast and memory-efficient alignment of short DNA sequences to the human genome.” Genome Biology 10: R25. 10.1186/gb-2009-10-3-r25.

Léran, S., K. H. Edel, M. Pervent, et al. 2015. “Nitrate sensing and uptake in *Arabidopsis* are enhanced by ABI2, a phosphatase inactivated by the stress hormone abscisic acid.” Science Signaling 8: ra43. DOI: 10.1126/scisignal.aaa4829.

Liu, K. H., C. Y. Huang, and Y. F. Tsay. 1999. “CHL1 is a dual-affinity nitrate transporter of Arabidopsis involved in multiple phases of nitrate uptake.” Plant Cell 11: 865–874. 10.1105/tpc.11.5.865.

Liu, K. H., Y. Niu, M. Konishi, et al. 2017. “Discovery of nitrate–CPK–NLP signalling in central nutrient–growth networks.” Nature 545: 311–316. 10.1038/nature22077.

Liu, K. H., M. Liu, Z. Lin, et al. 2022. “NIN-like protein 7 transcription factor is a plant nitrate sensor.” Science 377: 1419–1425. 10.1126/science.add1104.

Maeda, Y., M. Konishi, T. Kiba, et al. 2018. “A NIGT1-centred transcriptional cascade regulates nitrate signalling and incorporates phosphorus starvation signals in *Arabidopsis*.” Nature Communications 9: 1376. 10.1038/s41467-018-03832-6.

Marchive, C., F. Roudier, L. Castaings, et al. 2013. “Nuclear retention of the transcription factor NLP7 orchestrates the early response to nitrate in plants.” Nature communications 4: 1713. 10.1038/ncomms2650.

Medici, A., and G. Krouk. 2014. “The primary nitrate response: a multifaceted signalling pathway.” Journal of Experimental Botany 65: 5567–5576. 10.1093/jxb/eru245.

Miyawaki, K., M. Matsumoto-Kitano, and T. Kakimoto. 2004. “Expression of cytokinin biosynthetic isopentenyltransferase genes in *Arabidopsis*: tissue specificity and regulation by auxin, cytokinin, and nitrate.” Plant Journal 37: 128–138. 10.1046/j.1365-313X.2003.01945.x.

Miyawaki, K., P. Tarkowski, M. Matsumoto-Kitano, et al. 2006. “Roles of *Arabidopsis* ATP/ADP isopentenyltransferases and tRNA isopentenyltransferases in cytokinin biosynthesis.” Proceedings of the National Academy of Sciences of the United States of America 103: 16598–16603. 10.1073/pnas.0603522103.

Monden, K., T. Kamiya, D. Sugiura, T. Suzuki, T. Nakagawa, and T. Hachiya. 2022. “Root-specific activation of plasma membrane H^+^-ATPase 1 enhances plant growth and shoot accumulation of nutrient elements under nutrient-poor conditions in *Arabidopsis thaliana*.” Biochemical and Biophysical Research Communications 621: 39–45. 10.1016/j.bbrc.2022.06.097.

Nacry, P., E. Bouguyon, and A. Gojon. 2013. “Nitrogen acquisition by roots: physiological and developmental mechanisms ensuring plant adaptation to a fluctuating resource.” Plant and Soil 370: 1–29. 10.1007/s11104-013-1645-9.

Ohkubo, Y., M. Tanaka, R. Tabata, M. Ogawa-Ohnishi, and Y. Matsubayashi. 2017. “Shoot-to-root mobile polypeptides involved in systemic regulation of nitrogen acquisition.” Nature Plants 3: 17029. 10.1038/nplants.2017.29.

Okamoto, Y., T. Suzuki, D. Sugiura, T. Kiba, H. Sakakibara, and T. Hachiya. 2019. “Shoot nitrate underlies a perception of nitrogen satiety to trigger local and systemic signaling cascades in *Arabidopsis thaliana*.” Soil Science and Plant Nutrition 65: 56–64. 10.1080/00380768.2018.1537643.

Osugi, A., M. Kojima, Y. Takebayashi, N. Ueda, T. Kiba, and H. Sakakibara. 2017. “Systemic transport of *trans*-zeatin and its precursor have differing roles in *Arabidopsis* shoots.” Nature Plants 3: 17112. 10.1038/nplants.2017.112.

Ota, R., Y. Ohkubo, Y. Yamashita, M. Ogawa-Ohnishi, and Y. Matsubayashi. 2020. “Shoot-to-root mobile CEPD-like 2 integrates shoot nitrogen status to systemically regulate nitrate uptake in *Arabidopsis*.” Nature Communications 11: 641. 10.1038/s41467-020-14440-8.

Patterson, K., L. A. Walters, A. M. Cooper, et al. 2016. “Nitrate-regulated glutaredoxins control Arabidopsis primary root growth.” Plant Physiology 170: 989–999. 10.1104/pp.15.01776.

Poitout, A., A. Crabos, I. Petřík, et al. 2018. “Responses to systemic nitrogen signaling in Arabidopsis roots involve *trans*-zeatin in shoots.” Plant Cell 30: 1243–1257. 10.1105/tpc.18.00011.

Rashotte, A. M., L. R. Goertzen. 2010. “The CRF domain defines cytokinin response factor proteins in plants.” BMC Plant Biology 10: 74. 10.1186/1471-2229-10-74.

Remans, T., P. Nacry, M. Pervent, et al. 2006. “The *Arabidopsis* NRT1.1 transporter participates in the signaling pathway triggering root colonization of nitrate-rich patches.” Proceedings of the National Academy of Sciences of the United States of America 103: 19206–19211. 10.1073/pnas.060527510.

Riveras, E., J. M. Alvarez, E. A. Vidal, C. Oses, A. Vega, and R. A. Gutiérrez. 2015. “The calcium ion is a second messenger in the nitrate signaling pathway of Arabidopsis.” Plant Physiology 169: 1397–1404. 10.1104/pp.15.00961.

Rubin, G., T. Tohge, F. Matsuda, K. Saito, and W. R. Scheible. 2009. “Members of the *LBD* family of transcription factors repress anthocyanin synthesis and affect additional nitrogen responses in *Arabidopsis*.” Plant Cell 21: 3567–3584. 10.1105/tpc.109.067041.

Ruffel, S., G. Krouk, D. Ristova, D. Shasha, K. D. Birnbaum, and G. M. Coruzzi. 2011. “Nitrogen economics of root foraging: transitive closure of the nitrate–cytokinin relay and distinct systemic signaling for N supply vs. demand.” Proceedings of the National Academy of Sciences of the United States of America 108: 18524–18529. 10.1073/pnas.1108684108.

Sakakibara, H., K. Takei, and N. Hirose. 2006. “Interactions between nitrogen and cytokinin in the regulation of metabolism and development.” Trends in Plant Science 11: 440–448. 10.1016/j.tplants.2006.07.004.

Sakakibara, H. 2021. “Cytokinin biosynthesis and transport for systemic nitrogen signaling.” Plant Journal 105: 421–430. 10.1111/tpj.15011.

Safi, A., A. Medici, W. Szponarski, et al. 2021. “GARP transcription factors repress Arabidopsis nitrogen starvation response via ROS-dependent and -independent pathways.” Journal of Experimental Botany 72: 3881–3901. 10.1093/jxb/erab114.

Scheible, W. R., M. Lauerer, E. D. Schulze, M. Caboche, and M. Stitt. 1997. “Accumulation of nitrate in the shoot acts as a signal to regulate shoot-root allocation in tobacco.” Plant Journal 11: 671–691. 10.1046/j.1365-313X.1997.11040671.x.

Signora, L., I. De Smet, C. H. Foyer, and H. Zhang. 2001. “ABA plays a central role in mediating the regulatory effects of nitrate on root branching in *Arabidopsis*.” Plant Journal 28: 655–662. 10.1046/j.1365-313x.2001.01185.x.

Su, H., T. Wang, C. Ju, et al. 2021. “Abscisic acid signaling negatively regulates nitrate uptake via phosphorylation of NRT1. 1 by SnRK2s in *Arabidopsis*.” Journal of Integrative Plant Biology 63: 597–610. 10.1111/jipb.13057.

Tabata, R., K. Sumida, T. Yoshii, K. Ohyama, H. Shinohara, and Y. Matsubayashi. 2014. “Perception of root-derived peptides by shoot LRR-RKs mediates systemic N-demand signaling.” Science 346: 343–346. DOI: 10.1126/science.1257800.

Takei, K., N. Ueda, K. Aoki, et al. 2004a. “*AtIPT3* is a key determinant of nitrate-dependent cytokinin biosynthesis in *Arabidopsis*.” Plant & Cell Physiology 45: 1053–1062. 10.1093/pcp/pch119.

Takei, K., T. Yamaya, and H. Sakakibara. 2004b. “Arabidopsis *CYP735A1* and *CYP735A2* encode cytokinin hydroxylases that catalyze the biosynthesis of *trans*-zeatin.” Journal of Biological Chemistry 279: 41866–41872. 10.1074/jbc.M406337200.

Tokunaga, H., M. Kojima, T. Kuroha, et al. 2011. “Arabidopsis lonely guy (LOG) multiple mutants reveal a central role of the LOG-dependent pathway in cytokinin activation.” Plant Journal 69: 355–365. 10.1111/j.1365-313X.2011.04795.x.

Wang, R., R. Tischner, R. A. Gutiérrez, et al. 2004. “Genomic analysis of the nitrate response using a nitrate reductase-null mutant of Arabidopsis.” Plant Physiology 136: 2512–2522. 10.1104/pp.104.044610.

Wang, R, X. Xing, and N. M. Crawford. 2007. “Nitrite acts as a transcriptome signal at micromolar concentrations in Arabidopsis roots.” Plant Physiology 145: 1735–1745. 10.1104/pp.107.108944.

Wang, X., C. Feng, L. Tian, et al. 2021. “A transceptor–channel complex couples nitrate sensing to calcium signaling in *Arabidopsis*.” Molecular Plant 14: 774–786. 10.1016/j.molp.2021.02.005.

Werner, T., V. Motyka, V. Laucou, R. Smets, H. Van Onckelen, and T. Schmülling. 2003. “Cytokinin-deficient transgenic Arabidopsis plants show multiple developmental alterations indicating opposite functions of cytokinins in the regulation of shoot and root meristem activity.” Plant Cell 15: 2532–2550. 10.1105/tpc.014928.

Werner, T., K. Holst, Y. Pörs, et al. 2008. “Cytokinin deficiency causes distinct changes of sink and source parameters in tobacco shoots and roots.” Journal of Experimental Botany 59: 2659–2672. 10.1093/jxb/ern134.

Werner, T., Schmülling, T. 2009. “Cytokinin action in plant development.” Current Opinion in Plant Biology 12: 527–538. 10.1016/j.pbi.2009.07.002.

Wu, B., J. Meng, H. Liu, et al. 2023. “Suppressing a phosphohydrolase of cytokinin nucleotide enhances grain yield in rice.” Nature Genetics 55: 1381–1389. 10.1038/s41588-023-01454-3.

Xu, N., R. Wang, L. Zhao, et al. 2016. “The Arabidopsis NRG2 protein mediates nitrate signaling and interacts with and regulates key nitrate regulators.” Plant Cell 28: 485–504. 10.1105/tpc.15.00567.

Zhou, Y., B. Zhou, L. Pache, et al. 2019. “Metascape provides a biologist-oriented resource for the analysis of systems-level datasets.” Nature Communications 10: 1523. 10.1038/s41467-019-09234-6.

Zhang, H., A. Jennings, P. W. Barlow, and B. G. Forde. 1999. “Dual pathways for regulation of root branching by nitrate.” Proceedings of the National Academy of Sciences of the United States of America 96: 6529–6534. 10.1073/pnas.96.11.6529.

