## Supplemental Data 1 for "Shoot nitrate status regulates Arabidopsis shoot growth and systemic transcriptional responses via shoot adenosine phosphate-isopentenyltransferase 3"

### Figure S1

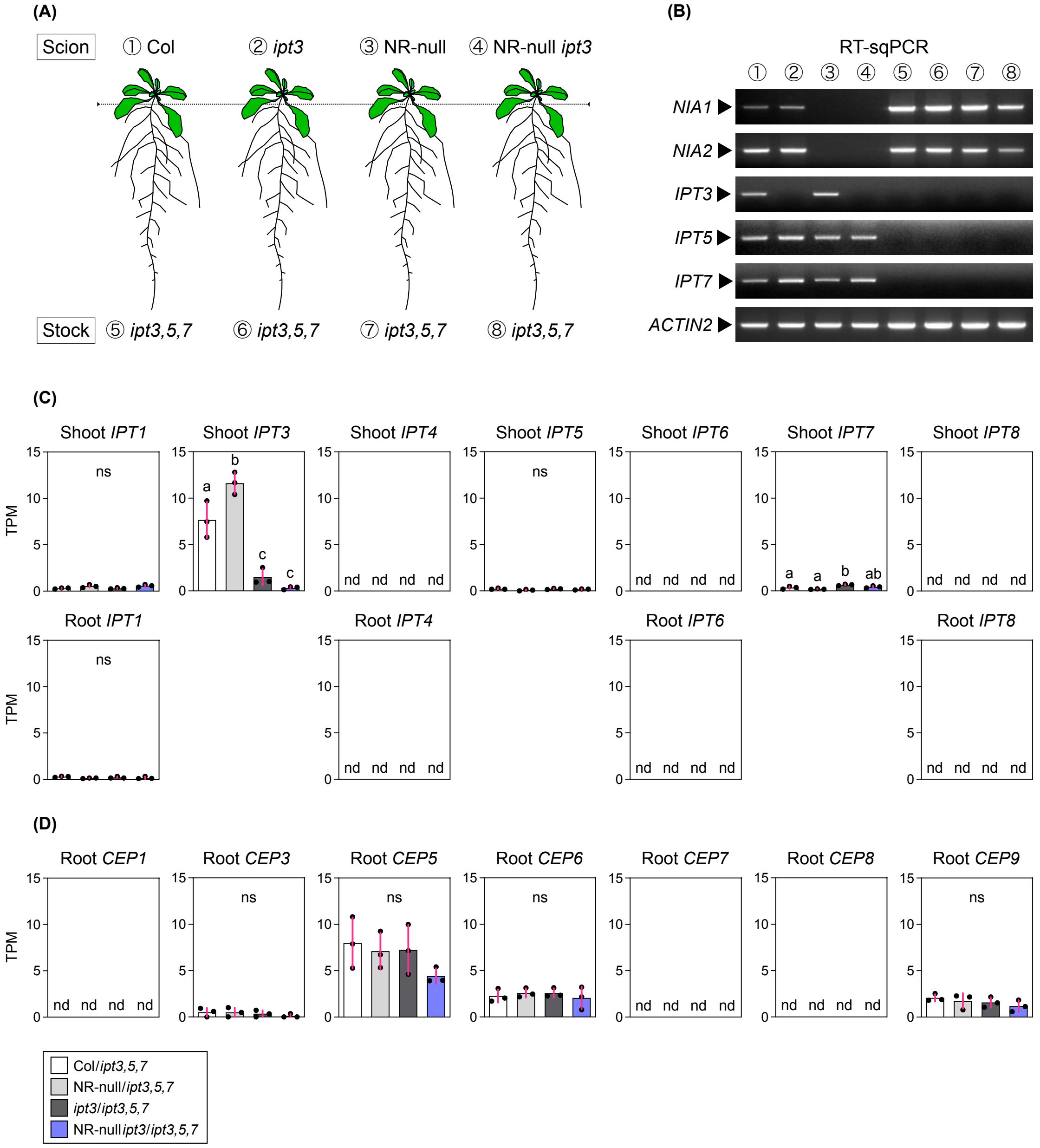


**Figure S1.** Confirmation of gene knockouts in the shoots and roots of grafted plants. (A) A schematic representation of four types of grafted plants. The dotted line represents the grafted seam in the hypocotyl. (B) A representative result of RT-sqPCR and agarose electrophoresis to confirm shoot and root expression of *NIA1*, *NIA2*, *IPT3*, *IPT5*, *IPT7*, and *ACTIN2* in grafted plants. The number of PCR cycles; 30 for *NIA1* and *NIA2*, 35 for *IPT3*, *IPT5*, *IPT7*, and 28 for *ACTIN2*. (C) TPM values of adenylate *IPT* genes as determined using RNA-seq in the shoots and roots of grafted plants 5 days after nitrogen removal (condition 5). (D) TPM values of nitrogen starvation-inducible *CEP* genes as determined using RNA-seq in the roots of grafted plants 5 days after nitrogen removal (condition 5).

### Figure S2

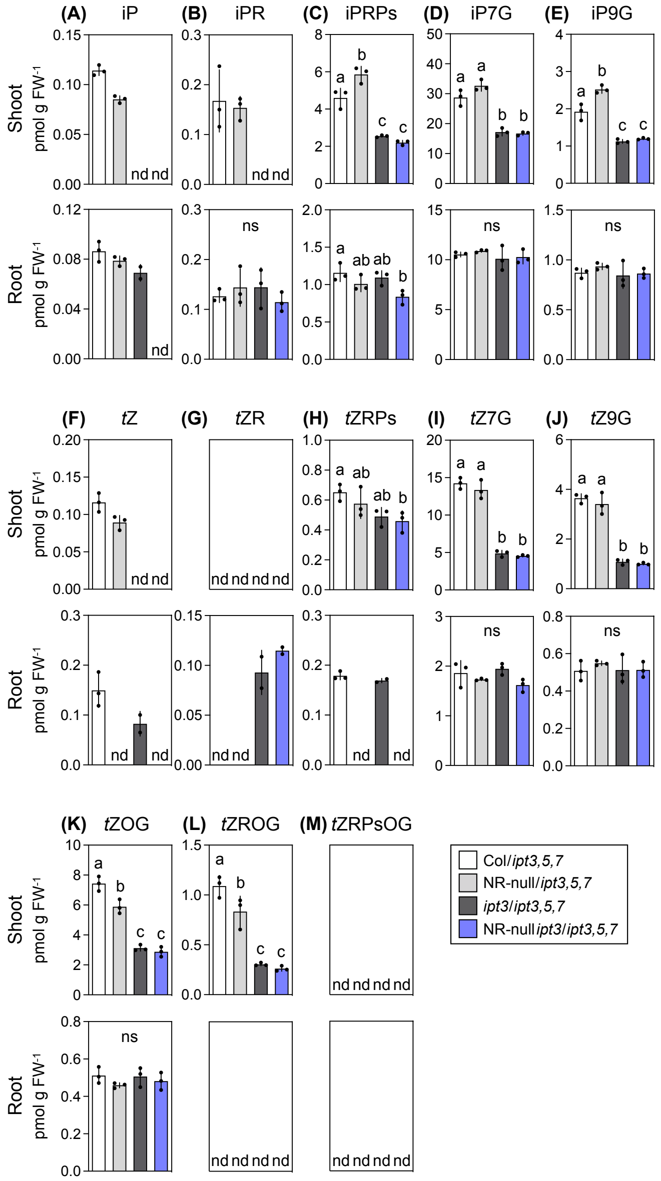


**Figure S2.** Effects of the shoot nitrate status and shoot *IPT3* expression on the shoot and root concentrations of CK species. (A–M) The shoot and root concentrations of iP (A), iPR (B), iPRPs (C), iP7G (D), iP9G (E), *t*Z (F), *t*ZR (G), *t*ZPRs (H), *t*Z7G (I), *t*Z9G (J), *t*ZOG (K), *t*ZROG (L), and *t*ZRPsOG (M) in the grafted plants 5 days after nitrogen removal (condition 5). Sampling was done once. In the sampling, three grafted plants was pooled as one biological replicate. Data are presented as mean ± SD [n = 3, excluding iP, *t*Z, *t*ZR, and *t*ZRPs concentrations in *ipt3*/*ipt3,5,7* roots and *t*ZR concentrations in NR-null *ipt3*/*ipt3,5,7* roots (n = 2)]. Different lowercase letters indicate significant differences, as determined via Tukey–Kramer tests ($P < 0.05$). iP, $N^6$-($\Delta^2$-isopentenyl)adenine; iPR, iP riboside; iPRPs, iP ribotides; iP7G, iP-7-*N*-glucoside; iP9G, iP-9-*N*-glucoside; *t*Z, *trans*-zeatin; *t*ZR, *t*Z riboside; *t*ZRPs, *t*Z ribotides; *t*Z7G, *t*Z-7-*N*-glucoside; *t*Z9G, *t*Z-9-*N*-glucoside; *t*ZOG, *t*Z-*O*-glucoside; *t*ZROG, *t*ZR-*O*-glucoside; *t*ZRPsOG, *t*ZRPs-*O*-glucoside; ns, not significant; nd, not detected; FW, fresh weight.

### Figure S3

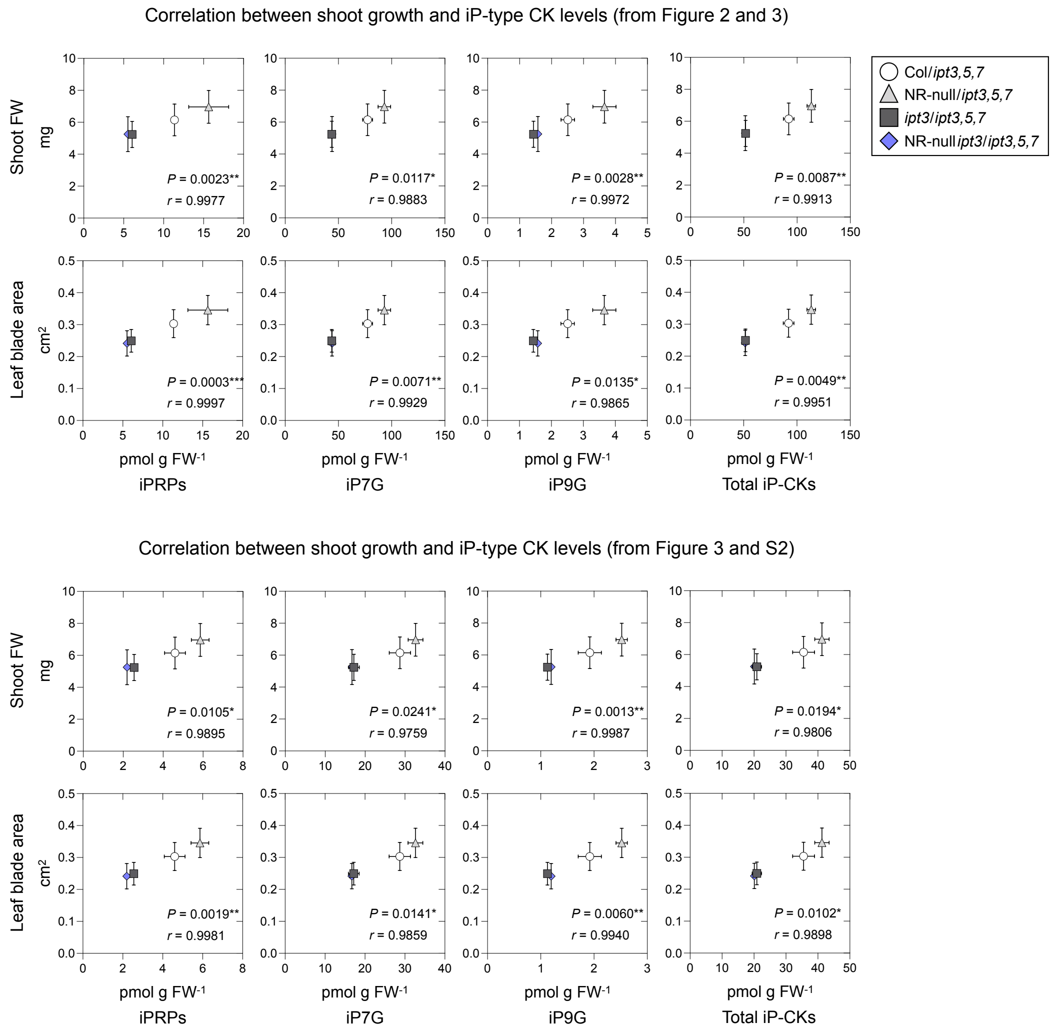


**Figure S3.** Correlation analysis between shoot growth and iP-type CK levels in the shoot. Statistical significance was conducted using Pearson correlation analysis (*$P$ < 0.05; **$P$ < 0.01; ***$P$ < 0.001). $r$ denotes Pearson's correlation coefficient.

### Figure S4

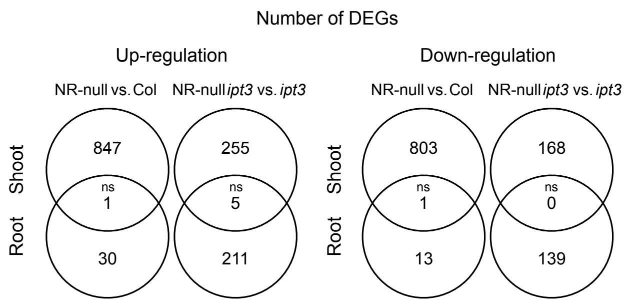


**Figure S4.** Venn diagram presenting the number of DEGs at a minimum fold change of 1.5 in shoots and roots. Statistical significance of overlap was assessed using the hypergeometric test ($P < 0.05$). ns, not significant.

### Figure S5

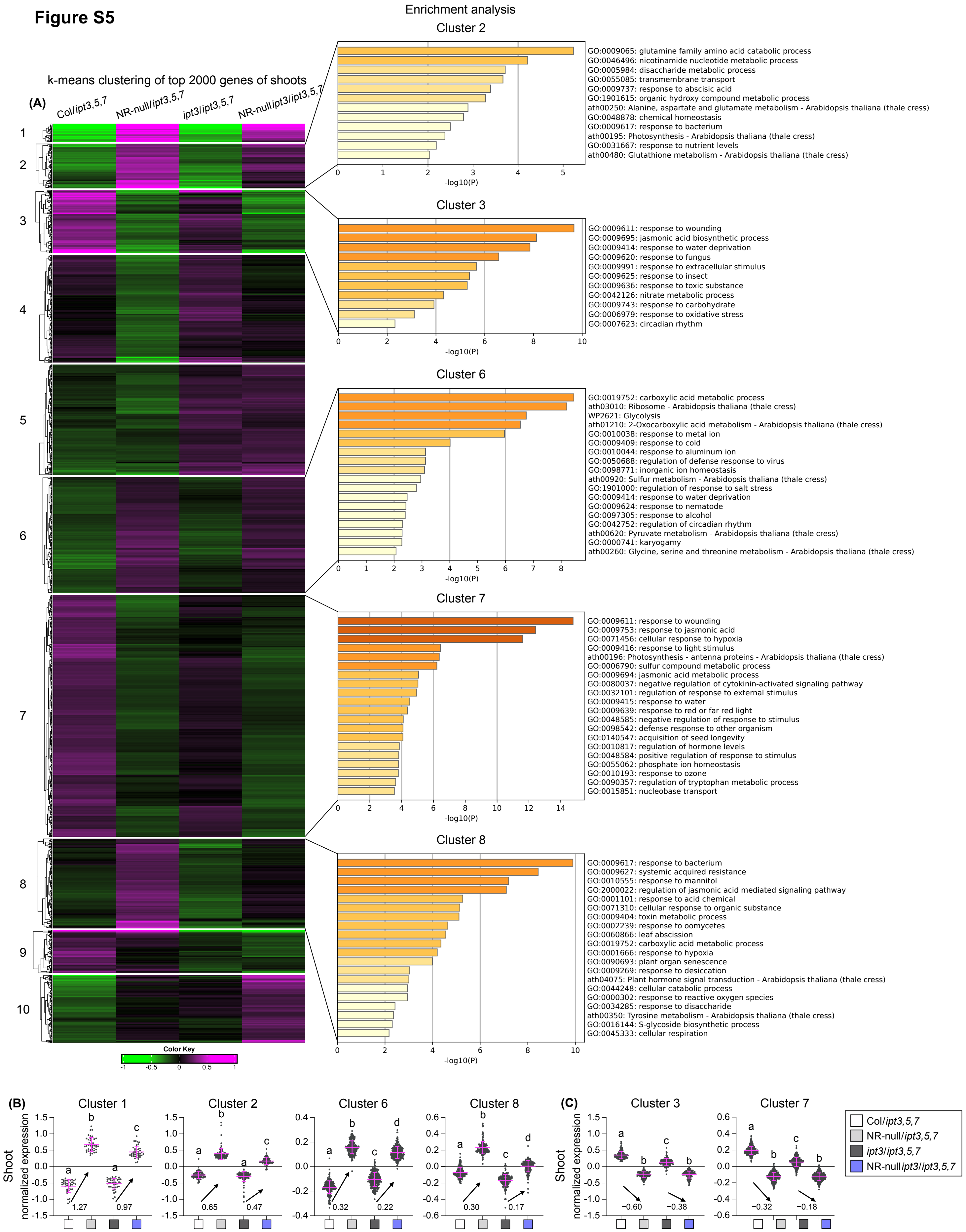


**Figure S5.** Classification of transcripts via k-means clustering in shoots of grafted plants 5 days after nitrogen removal (condition 5). (A) Heat map from k-means clustering of mean normalized transcript levels with outputs of enriched terms. The genes for which the standard deviation was ranked in the top 2000 were used for the clustering. Magenta and green represent higher and lower expression levels, respectively. B) Plots of mean normalized transcript levels of the genes upregulated by shoot NR deficiency, the responses of which were enhanced by shoot IPT3. (C) Plots of mean normalized transcript levels of the genes downregulated by shoot NR deficiency, the responses of which were enhanced by shoot IPT3. Different lowercase letters indicate significant differences, as determined via Tukey–Kramer test ($P < 0.05$).

### Figure S6

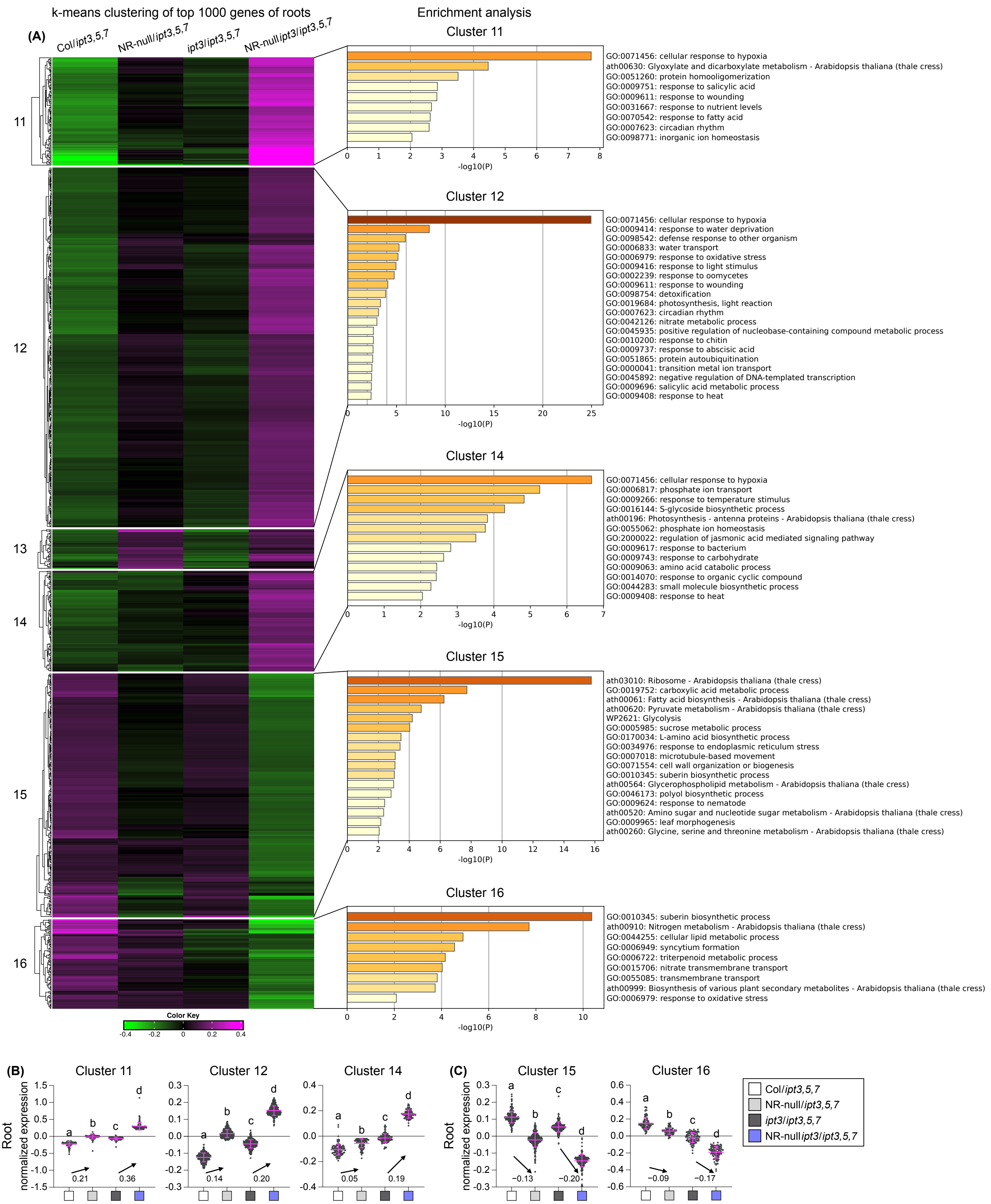


**Figure S6.** Classification of transcripts via k-means clustering in in roots of grafted plants 5 days after nitrogen removal (condition 5). (A) Heat map from k-means clustering of mean normalized transcript levels with outputs of enriched terms. The genes for which the standard deviation was ranked in the top 1000 were used for the clustering. Magenta and green represent higher and lower expression levels, respectively. (B) Plots of mean normalized transcript levels of the genes upregulated by shoot NR deficiency, the responses of which were dampened by shoot IPT3. (C) Plots of mean normalized transcript levels of the genes downregulated by shoot NR deficiency, the responses of which were dampened by shoot IPT3. Different lowercase letters indicate significant differences, as determined via Tukey–Kramer test ($P < 0.05$).

### Figure S7

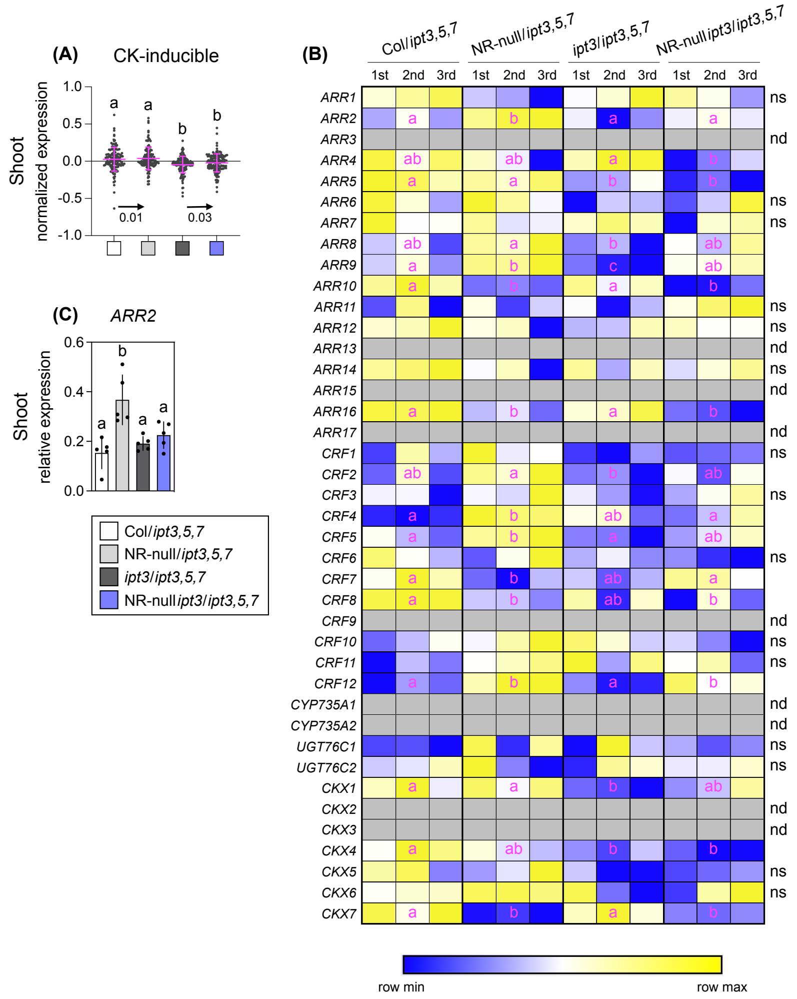

**Figure S7.** Dissection of CK-related genes in shoots of grafted plants 5 days after nitrogen removal (condition 5). (A) Plots of mean normalized transcript levels of the CK-inducible genes (mean ± SD). The gene list was obtained from Bhargava et al. (2013). (B) Transcriptional changes in genes involved in CK signaling/metabolism. The normalized transcript levels were visualized as heatmaps using MORPHEUS (https://software.broadinstitute.org/morpheus). The relative color scheme uses the minimum (blue) and maximum (yellow) values in each row. 1st, 2nd, and 3rd denote the order of three independent experiments. (C) Transcript levels of *ARR2*, as determined using RT-qPCR. Data are presented as mean ± SD (n = 5). (A–C) Different lowercase letters indicate significant differences, as determined via Tukey–Kramer tests at *P* < 0.05. ns, not significant; nd, not detected.

### Figure S8

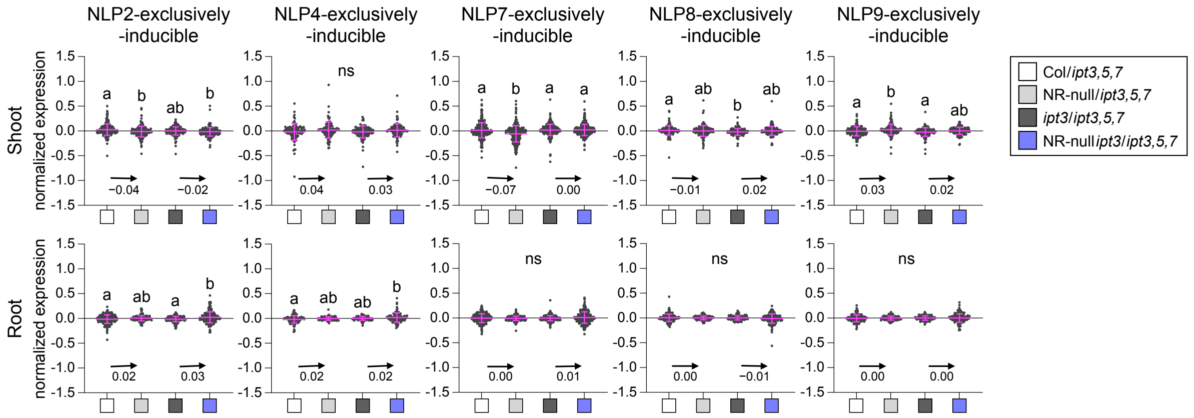


**Figure S8.** Plots of mean normalized transcript levels of the genes induced exclusively by NLP2, NLP4, NLP7, NLP8, or NLP9 in shoots and roots of grafted plants 5 days after nitrogen removal (condition 5). Different lowercase letters indicate significant differences, as determined via Tukey–Kramer tests at $P < 0.05$. The gene list was obtained from Liu et al. (2022). ns, not significant.

### Figure S9

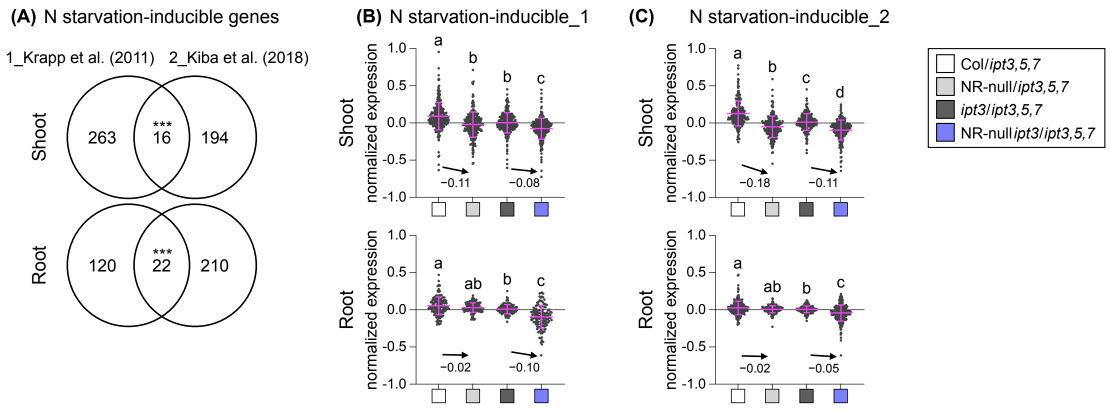


**Figure S9.** Effects of the shoot nitrate status and shoot *IPT3* expression on nitrogen starvation-inducible genes in shoots and roots of grafted plants 5 days after nitrogen removal (condition 5). (A) Venn diagram presenting the number of nitrogen starvation-inducible genes in Krapp et al. (2011) and Kiba et al. (2018). Statistical significance of overlap was assessed using the hypergeometric test (**P* < 0.05; ***P* < 0.01; ****P* < 0.001). (B, C) Plots of mean normalized transcript levels of nitrogen starvation-inducible genes. The gene lists were obtained from Krapp et al. (2011) for (B) and Kiba et al. (2018) for (C). Different lowercase letters indicate significant differences, as determined via Tukey–Kramer test (*P* < 0.05). The numbers on the graph represent the difference in mean values.

### Figure S10

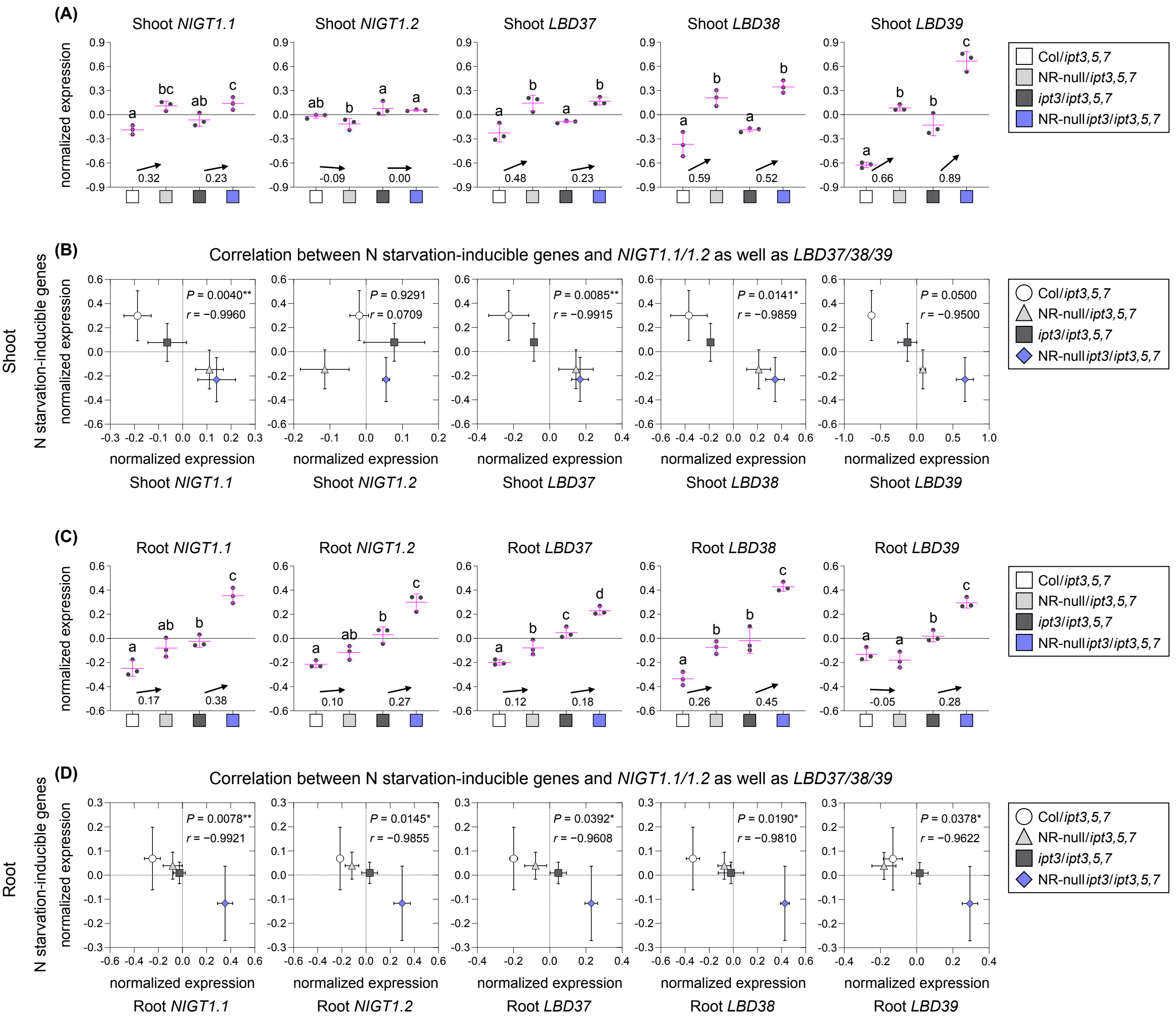

**Figure S10.** (A, C) Plots of normalized transcript levels of repressor genes in shoots (A) and roots (C) of grafted plants 5 days after nitrogen removal (condition 5). (B, D) Correlation analysis between the mean normalized transcript levels of nitrogen starvation-inducible genes and repressor genes in shoots (B) and roots (D). Statistical significance was conducted using Pearson correlation analysis (**P* < 0.05; ***P* < 0.01; ****P* < 0.001). *r* denotes Pearson's correlation coefficient.

### Figure S11

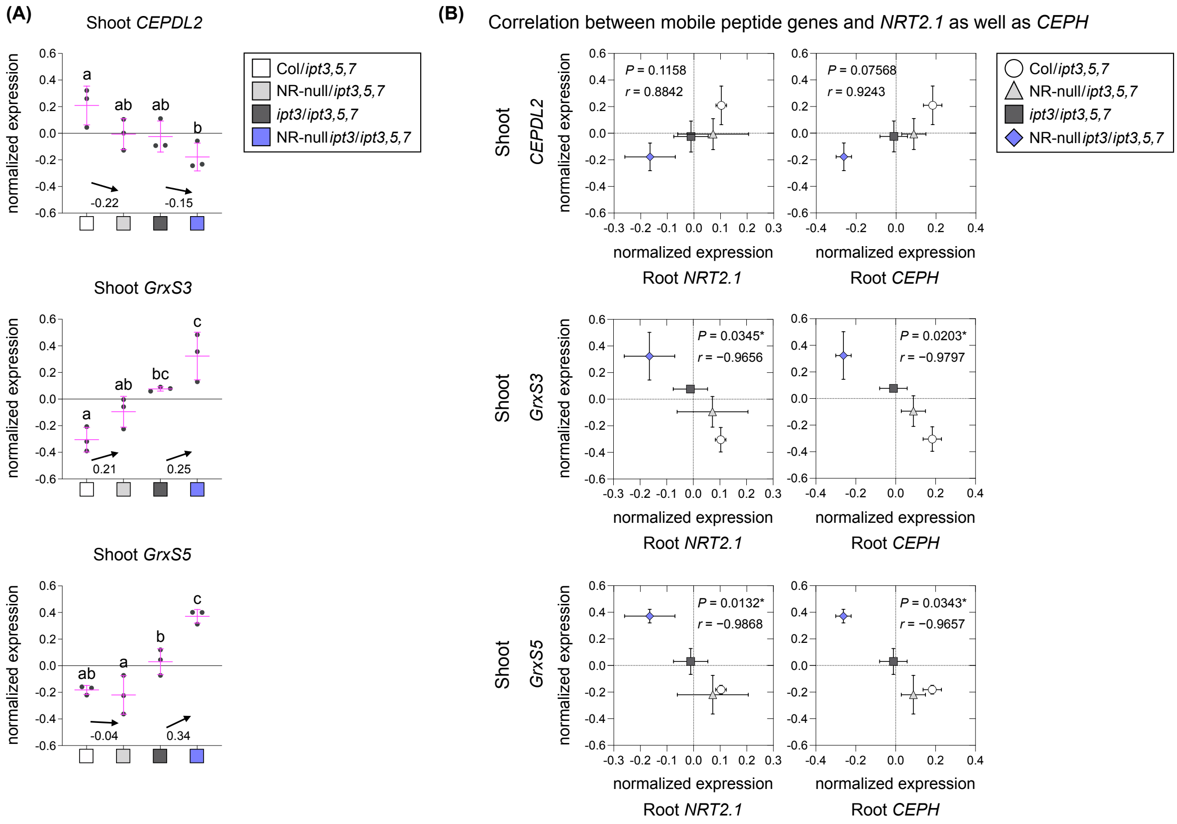


**Figure S11.** (A) Plots of normalized transcript levels of shoot-to-root mobile peptide genes in shoots of grafted plants 5 days after nitrogen removal (condition 5). (B) Correlation analysis between the mean normalized transcript levels of shoot-to-root mobile peptide genes in shoots and genes involved in high-affinity nitrate transport in roots. Statistical significance was conducted using Pearson correlation analysis (**P* < 0.05; ***P* < 0.01; ****P* < 0.001). *r* denotes Pearson's correlation coefficient.
